# A cryptic START domain regulates deeply conserved transcription factors

**DOI:** 10.1101/2025.07.29.667167

**Authors:** Courtney E. Dresden, Ekaterina P. Andrianova, Brian J. Smith, Nicole I. Callery, Dominic Kolonay, Ashton S. Holub, Ricardo Urquidi Camacho, Sarah G. Choudury, Isabella J. Higgins, Igor B. Zhulin, Aman Y. Husbands

**Affiliations:** Department of Biology, University of Pennsylvania, 415 S. University Ave, Philadelphia, PA, 19104, USA; Molecular, Cellular, and Developmental Biology, the Ohio State University, Columbus, OH 43215, USA; Department of Microbiology, The Ohio State University, Columbus, OH 43215, USA; Department of Molecular Genetics, The Ohio State University, Columbus, OH 43215, USA; Epigenetics Institute, University of Pennsylvania, Philadelphia PA 19104 USA

## Abstract

Transcription factors (TFs) integrate a diverse array of inputs to achieve the exquisite control of gene expression necessary for life. In plants, this is exemplified by the deeply conserved CLASS III HOMEODOMAIN LEUCINE ZIPPER (HD-ZIPIII) family of TFs. HD-ZIPIII activity is controlled by inputs at transcriptional, post-transcriptional, and post-translational levels. As part of their multidomain architecture, HD-ZIPIII TFs contain a StAR-related lipid transfer (START) domain, a ubiquitously distributed evolutionary module that binds various types of lipophilic ligands. Here, we show that HD-ZIPIII and HD-ZIPIV proteins contain a cryptic, deeply conserved START domain which we term the disorder-containing START domain (dSTART). The dSTART domain is required for HD-ZIPIII developmental function, controlling their subcellular localization and DNA-binding properties. The dSTART domain also helps discriminate responsive from non-responsive binding sites across the HD-ZIPIII shared genetic network. Finally, we identify candidate ligands of the dSTART domain including several species of phosphatidylglycerol and phosphatidic acid. The identification and functional characterization of a cryptic START domain provides new mechanistic insights into a deeply conserved family of TFs with roles in nearly all aspects of plant development.

## INTRODUCTION

In biology, highly complex processes unfold in precise stereotypical ways [1]. This is accomplished in large part by transcription factors (TFs) which integrate numerous genetic and environmental inputs to coordinate gene expression [2]. To achieve the required precision, these important regulators are often controlled by multiple distinct mechanisms [3,4]. For instance, TFs are preferred targets of microRNAs in both plants and animals [5,6], and post-transcriptional regulation of TF activity is essential for organismal growth and development [7–9]. At the protein level, TF activity is modulated by numerous inputs including interacting partners, various types of post-translational modifications, and, in some cases, binding of small-molecule ligands [10–14]. Understanding how highly regulated TFs integrate a diverse array of inputs to produce stable transcriptional outcomes is a perennial challenge, and mapping the full suite of inputs converging on a given TF is an essential prerequisite.

In plants, an excellent example of highly regulated TFs are the members of the deeply conserved CLASS III HOMEODOMAIN LEUCINE ZIPPER (HD-ZIPIII) family. HD-ZIPIII proteins arose over 700 million years ago, sometime after the divergence of land plants from unicellular algae, then proliferated throughout evolution [15]. The HD-ZIPIII family in *Arabidopsis thaliana* has five paralogs *– REVOLUTA (REV), PHABULOSA (PHB), PHAVOLUTA (PHV), CORONA (CNA),* and *ATHB-8* – that collectively impact nearly all aspects of development through functionally redundant and functionally divergent activities. Examples of redundancy include *PHB, PHV, REV,* and *CNA* promoting dorsal identity in lateral organs and xylem cell fate in the vasculature [16]. An example of functional divergence is *REV* and *CNA*, whose activities promote and antagonize the stem cell niche, respectively [16]. In keeping with their critical roles in development, HD-ZIPIII activity is regulated by the integration of multiple inputs. One such input is microRNA 166 (miR166), which spatially restricts HD-ZIPIII transcript accumulation [17,18]. Loss of miR166 regulation leads to a wide range of pleiotropic phenotypes including enlarged meristems, shortened roots, mispatterned vasculature, dorsalized leaves, and floral patterning defects [19–24]. Other inputs into HD-ZIPIII activity include interacting proteins from several TF and chromatin remodeler classes [25–28], as well as members of the LITTLE ZIPPER (ZPR) family of microProteins [29–31]. The latter are directly transactivated by HD-ZIPIII TFs and negatively regulate HD-ZIPIII activity through the formation of heteromeric complexes lacking DNA-binding capacity [29,30].

HD-ZIPIII TFs contain a DNA binding homeodomain (HD), a leucine zipper dimerization domain (LZ), and a C-terminal PAS-like MEKHLA domain [31,32]. These proteins also contain a StAR-related transfer (START) domain suggesting additional levels of regulation. START domains are part of the StARkin domain superfamily, named for their kinship to the steroidogenic acute regulatory (StAR) protein [33]. StARkin domains are present throughout the tree of life and are defined by a conserved α/β helix grip fold structure which binds a variety of ligands, which include sterols, phospholipids, and isoprenoids [34–36]. Ligand binding triggers conformational changes which often directly modulate the activity of StARkin-containing multidomain proteins via a broad set of regulatory mechanisms [37]. The presence of a ligand-binding domain in HD-ZIPIII proteins led to the long-standing hypothesis that their activity is controlled by lipid ligands, perhaps analogous to nuclear receptors (NRs) in metazoan systems [15,37].

Supporting this, several types of phospholipids were recently identified as ligands for the HD-ZIPIII START domain including multiple species of phosphatidylcholine (PC). Mutations that prevent PC-binding abolish DNA-binding competence and hence HD-ZIPIII transcriptional output [38]. The START domain also promotes HD-ZIPIII homodimerization and increases transcriptional potency. These findings propose a model in which the START domain converts HD-ZIPIII proteins into transcriptionally potent, DNA-binding-competent TFs upon binding of their phospholipid ligand [38]. START-dependent effects on homodimerization have also been reported for the evolutionarily related HD-ZIPIV family of TFs [39]. However, the HD-ZIPIV START domain does not seem to affect DNA-binding competence [39], instead impacting HD-ZIPIV activity through modulation of subcellular localization and protein stability by ceramide and lyso-PC ligands [40,41]. These findings exemplify the flexibility of the StARkin module, as even evolutionarily related sets of proteins can use dramatically different regulatory mechanisms.

The START domain also contributes to functional divergence of HD-ZIPIII paralogs. A recent study found that HD-ZIPIII TFs do not generate their paralog-specific transcriptional outcomes by binding to distinct sites in the genome [42]. Instead, functional divergence results from differential usage of shared binding sites, with hundreds of uniquely regulated genes emerging from a commonly bound genetic network. After binding to a given site, HD-ZIPIII TFs appear to independently determine whether this will lead to changes in gene expression. If the site is considered ‘responsive’ for the paralog, then transcription is either promoted or inhibited. If the site is considered ‘non-responsive’ for the paralog, then gene expression remains unchanged. Discrimination between responsive and non-responsive binding sites depends in part on the START domain, however the regulatory properties driving this discrimination are unknown [42].

Downstream of the HD-ZIPIII START domain is a region with no known structural features. Here, we show that this region contains a cryptic START domain. This domain is present in all HD-ZIPIII proteins, as well as members of the HD-ZIPIV subfamily, indicating a deep ancestry. Subsequent predictive modeling of HD-ZIPIII and HD-ZIPIV multimers using AlphaFold2 similarly recognizes this cryptic domain ([43]; see Discussion). In addition to amino acids and structural features common to StARkin domains, the newly identified START domain possesses unique defining features. These include a highly conserved LPSGF motif as well as long insertions between three of its secondary structure elements. One of these insertions has properties of an intrinsically disordered region (IDR) prompting us to name this new domain the disorder-containing START domain (dSTART) to highlight the unique feature that allowed it to evade detection by sequence-based algorithms for over twenty years. Genetic and molecular analyses using PHB as a representative HD-ZIPIII TF reveal the dSTART domain is required for HD-ZIPIII developmental function, and controls subcellular localization and DNA-binding competence and/or affinity. In addition, genomic assays propose a role for the dSTART domain in helping HD-ZIPIII TFs discriminate between responsive and non-responsive binding sites across their shared genetic network. Finally, we identify high-confidence candidate ligands of the dSTART domain belonging to two separate, but structurally related, phospholipid classes. Our findings provide a wealth of new mechanistic insights into a deeply conserved, essential family of TFs with roles in nearly all aspects of plant development.

## RESULTS

### HD-ZIPIII TFs contain a cryptic START domain

Genetic analyses have long suggested a regulatory role for the region downstream of the HD-ZIPIII START domain [16]. This region is sometimes referred to as the HD-START-associated domain [44,45], however there was previously no evidence of any structural domain(s) (**Fig. 1a**). This has hindered the molecular identification of new HD-ZIPIII regulatory inputs, prompting us to examine it more carefully for structural features that may have been missed by previous analyses. All *Arabidopsis* HD-ZIPIII paralogs (REVOLUTA (REV), PHABULOSA (PHB), PHAVOLUTA (PHV), CORONA (CNA), and ATHB8) have similar architectures, and we selected PHB as a representative HD-ZIPIII protein. Initial analyses confirm the four established domains of PHB: HD (Pfam ID PF00046), LZ (or bZIP_1; Pfam ID PF00170), START (Pfam ID PF01852), and MEKHLA (Pfam ID PF08670). By contrast, no domain was recognized within the ∼250 amino acid sequence separating the START and MEKHLA domains (residues 385 to 702). Searches initiated with this sequence against the Pfam and Conserved Domain Databases (CDD) also found no similarity to any known domain. We then conducted a more sensitive profile-profile matching search using HHpred [46]. HHpred analyses identified several proteins from *Mus musculus* and *Homo sapiens* with high confidence support (>90% probability; PDB IDs of five best hits: 2PSO,2MOU, 6L1M, 5I9J, 2R55; **Fig. 1a**), all of which contain a StARkin domain from the START subfamily (Pfam ID PF01852).

**Figure 1.**
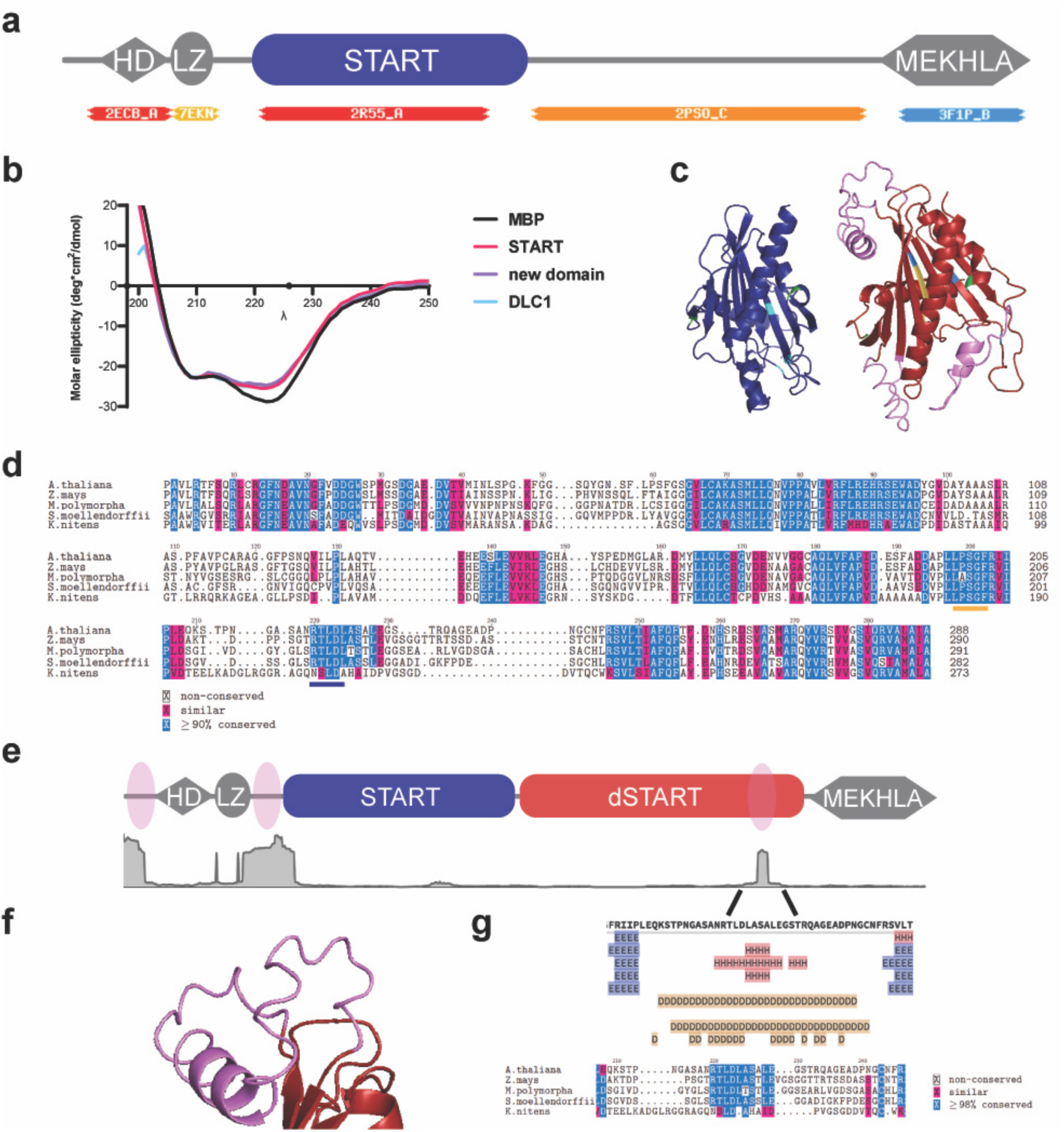
HD-ZIPIII proteins contain a cryptic, deeply conserved START domain defined by unique structural motifs and features. **a** Classically defined HD-ZIPIII protein architecture (HD = homeodomain, LZ = leucine zipper, START = StAR-related lipid transfer, MEKHLA/PAS). HHpred analyses show the region between the START and MEKHLA domains has homology to the StARkin protein STARD6 (PDB ID: 2MOU_A) while the START domain resembles human STARD5 (PDB ID: 2R55). **b**, Circular dichroism of MBP-START, MBP-new domain, and MBP-DLC1 shows all three recombinant proteins have virtually identical secondary structures. **c**, Homology-guided modeling shows the PHB START (blue) and new domain (red) domains each adopt the helix-grip fold structure characteristic of StARkin domains. Regions of interest include highly conserved tryptophan (green) and arginine or glutamine residues (blue), an LPSGF motif (yellow), and three largely unstructured loops or insertions (pink). **d**, Minimal multiple sequence alignment reveals deep conservation of the dSTART domain. Regions of interest include the LPSGF (yellow bar) and RTLDLA (blue bar) motifs. **e**, PHB architecture (upper) with DEPICTER readout (lower; gray graph). Two regions of intrinsic disorder flank the HD and LZ, and a third region localizes near the C-terminus of the new disorder-containing START (dSTART) domain. **f**, AlphaFold2 homology model of the dSTART domain focusing on region of disorder predicted by DEPICTER (pink). **g**, Quick2D analysis indicates this region is predicted as both ordered and disordered (upper). Predictions of order occur at an RTLDLASXLE amino acid motif. MSAs indicate a deep conservation of this motif (lower).

We next built a multiple sequence alignment (MSA) comparing the newly discovered domain (residues 399 to 681) with the PHB START domain (residues 168 to 384) and the mammalian START domains returned by HHpred (**Supp Fig. 1**). Consistent with MSAs of other distantly related StARkin domains [47], relatively few amino acids are universally conserved. Importantly, these conserved residues include two tryptophans common to nearly all START domains [48], that are proposed to be important for structural stability or lipid ligand loading [49,50]. Mapping of secondary structure elements onto the MSA shows good agreement between the newly discovered domain and both mammalian and plant START sequences (**Supp Fig. 1**). To experimentally validate this prediction, we produced recombinant protein in *E. coli* and performed circular dichroism. These analyses confirm the new domain adopts virtually identical secondary structures as experimentally-validated START domains from PHB and STARD12 (Deleted in Liver Cancer-1 (DLC1); [28]; **Fig. 1b**). Finally, homology modeling with iTASSER [51–53] – and later AlphaFold2 [54] – predicts this new domain adopts the helix-grip fold structure characteristic of StARkin domains (**Fig. 1c**). Taken together, our computational and experimental data demonstrate that the uncharacterized region between the START domain and MEHKLA domain of HD-ZIPIII proteins contains a cryptic START domain.

### The cryptic START domain is deeply conserved and has unique structural features

To determine the extent of its conservation and any potentially important structural features, we built an MSA of the new START domain using HD-ZIPIII sequences from across the plant lineage. The domain is present in *Chlorokybus atmophyticus*, the earliest-diverging organism known to contain an HD-ZIPIII protein, as well as in members of the evolutionarily related HD-ZIPIV subfamily, indicating a deep ancestry that predates the divergence of the two HD-ZIP subfamilies (**Figs 1d; Supp Figs. 2, 3,** [43]). Our MSAs also reveal structural features unique to the new START domain. First is a five amino acid motif – LPSGF – which is absolutely conserved in >90% of the 2800 protein sequences included in our analysis (**Fig. 1d**; **Supp Doc 1**; see **Methods**). These residues partially overlap the seventh β-strand and are predicted to line the interior of the hydrophobic START binding pocket (**Fig. 1c**). The LPSGF motif is not present in other START domains and thus appears to be a major determinant of this new subfamily. Second, the new START domain is 30 to 80 amino acids larger than other described START domains. In HD-ZIPIII proteins, these additional residues manifest as three large insertions between the β1-β2, β7-β8, and α3-β2 secondary structure elements (**Fig. 1c; Supp Fig. 1**). By contrast, HD-ZIPIV proteins only possess the α3-β2 insertion (**Supp Fig. 4**). These insertions are likely the reason this new START domain was not recognized by previous generations of domain discovery tools such as Pfam or CDD. Interestingly, the α3-β2 insertion is predicted to be an intrinsically disordered region (IDR; **Figs 1e; Supp Fig. 4**). We thus renamed the cryptic START domain *disorder-containing START* (*dSTART*) to highlight the unique feature that allowed it to evade detection by sequence-based algorithms for over twenty years.

The internal IDR of the HD-ZIPIII dSTART domain has additional properties of interest. For instance, despite its prediction as an IDR, homology models reveal a small, weakly-supported α-helix (**Fig. 1f**). To investigate this possibility, the PHB dSTART sequence was run through Quick2D which integrates results from secondary structure, transmembrane, and disorder prediction packages [55]. Interestingly, this package finds support for both intrinsic disorder and a small α-helix (RTLDLASALE) within the PHB dSTART IDR (**Fig. 1g**). This property is reminiscent of regions that undergo signal-dependent transitions between ordered and disordered states [56]. In addition, our MSA shows minimal conservation of the IDR across plant evolution, with the notable exception of the predicted α-helical region (**Fig. 1g**). Similar analyses with the HD-ZIPIV dSTART domain find only intrinsic disorder (**Supp Fig. 4**), suggesting the RTLDLASXLE α-helix may have HD-ZIPIII-specific regulatory roles. Taken together, our analyses identify a new deeply conserved START domain with structural features that distinguish it from other StARkin domains.

### START and dSTART domains have distinct patterns of evolutionary conservation

The START and dSTART domains are both present in the earliest-identified HD-ZIPIII protein indicating a deep conservation. To begin to map the evolutionary relationship of these two domains, we analyzed their patterns of conservation across the plant lineage. This was done by creating a series of percent identity matrices using START and dSTART domain sequences from HD-ZIPIII paralogs in *Arabidopsis*, HD-ZIPIII orthologs in other plant species, and StARkin domains from non-plant species. Analyses were seeded using the sequence of CNA and its orthologs, as recent analyses indicate the ancestral HD-ZIPIII gene most closely resembles CNA [42].

We first compared the *Arabidopsis* CNA sequence to its four paralogs, which fall into two evolutionary subclades: the REV clade (PHB, PHV, and REV), and the CNA clade (CNA and ATHB8) (**Supp Table 1a**) [16]. The CNA START domain is relatively well conserved between paralogs, ranging from 73% identity for PHB to 80% identity for ATHB8 (**Fig. 2a; Supp Fig. 5A; Supp Table 1a**). By contrast, conservation is weaker for the CNA dSTART domain, particularly for members of the REV clade, with CNA and REV dSTART domains sharing only 62% identity (**Fig. 2a; Supp Table 1a**). We hypothesized this weaker conservation may result from the three dSTART insertions, which are variable in both size and amino acid composition (**Supp Fig. 5B**). To test this possibility, we removed the insertions from dSTART sequences. This qualitatively improved MSA alignments and increased percent identities between the paralogs (**Fig. 2b; Supp Fig. 5C, Supp Table 1**). However, these identities remained lower than those of the START domain suggesting divergence of the dSTART domain across the HD-ZIPIII subclades is only partially explained by these structural features.

**Figure 2.**
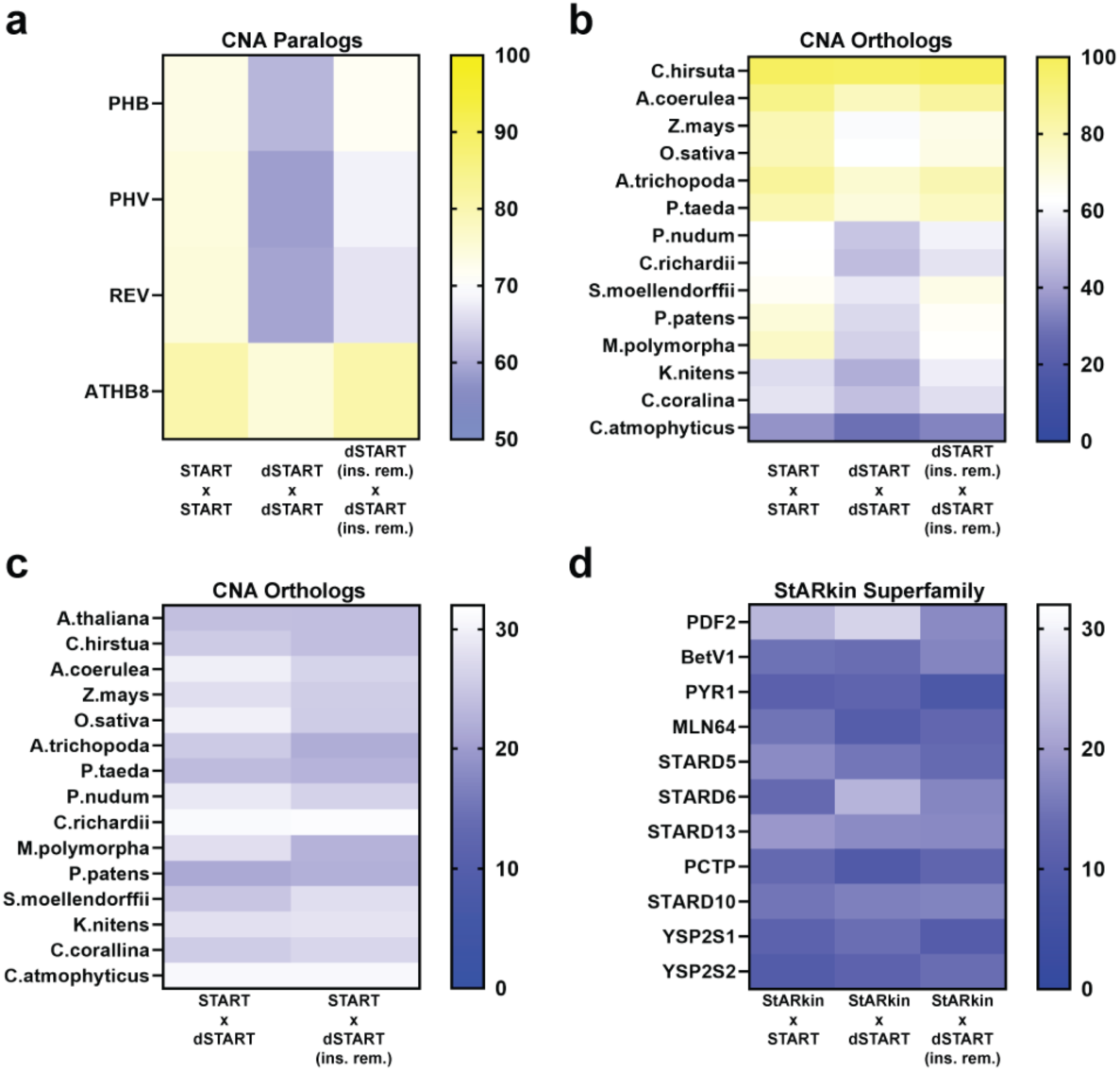
The START and dSTART domains have distinct patterns of evolutionary conservation. **a**, Percent identity matrices comparing START, dSTART, and dSTART (no insertions) sequences of CNA with its paralogs. The dSTART domain is less conserved than the START domain, and this is only partly explained by its multiple insertions. **b**, Percent identity matrices comparing START, dSTART, and dSTART (no insertions) sequences of CNA with its orthologs in extant plant species similarly indicate weaker conservation of the dSTART domain. **c**, Percent identity matrices comparing dSTART and dSTART (no insertions) sequences of CNA orthologs with their respective START domains. Percent identities of START-dSTART pairs range from 21%-31%, and removal of dSTART insertions usually lowers this conservation. **d**, Percent identity matrices comparing START, dSTART, and dSTART (no insertions) sequences of CNA with other members of the StARkin superfamily. Percent identities of START-dSTART pairs range from 10%-22% and removal of dSTART insertions has variable effects on this conservation.

We then determined the conservation patterns of the HD-ZIPIII START and dSTART domains across plant evolution by comparing CNA orthologs from several extant species (**Supp Table 1b**). Like the HD-ZIPIII paralogs, the START domain was more highly conserved than the dSTART domain in all ortholog comparisons (**Fig. 2b; Supp Table 1b)**. Conservation of dSTART between the orthologs was also increased by removing its insertions but rarely to levels seen with the START domain (**Fig. 2b; Supp Table 1b, Supp Fig. 6**). Thus, analyses with HD-ZIPIII paralogs and orthologs both support a weaker evolutionary conservation of the dSTART domain which is only partially explained by its insertions. Interestingly, this pattern is reversed in an HD-ZIPIV protein, whose dSTART domain consistently shows higher conservation than their START domain (**Supp Fig. 7; Supp Table 1c**).

Finally, we used these percent identity matrices to investigate potential evolutionary origins of the START and dSTART domains. For instance, their tandem arrangement could have arisen by capture of two distinct domains or by duplication of a single ancestral domain. Both possibilities are common mechanisms of domain architecture evolution [57]. To distinguish between these possibilities, we compared START-dSTART pairs within CNA orthologs of extant plant species. Depending on the species, a given CNA START domain shares 21%-31% (µ = 27%) identity with its adjacent dSTART domain (**Fig. 2c; Supp Table 1d**). By contrast, similar comparisons find only 10%-22% (µ = 14%) identity between other members of the StARkin superfamily and the CNA START or dSTART domain (**Fig. 2d; Supp Table 1e**). These findings support the idea of an ancient duplication, rather than capture of two distinct StARkin domains by the HD-ZIPIII or HD-ZIPIV antecedent.

Taken together, our evolutionary analyses reveal distinct patterns of START and dSTART domain conservation within and between the HD-ZIP subfamilies. In addition, the insertions that define the dSTART domain contribute to, but do not fully explain, the increased divergence of the dSTART domain in HD-ZIPIII proteins.

### Highly conserved StARkin residues within the dSTART domain are required for full PHB function

Next, we assessed dSTART-dependent effects on HD-ZIPIII developmental and molecular function using PHB as a representative member. We first tested the impact of the two tryptophan residues that are highly conserved across the StARkin superfamily by mutagenizing them to alanines (dWmut; **Fig. 1c**; [48]). These mutations are predicted to minimally affect protein folding (**Supp Fig. 8a**), and circular dichroism confirmed wildtype and dWmut dSTART domains purified from *E. coli* adopt virtually identical secondary structures (**Supp Fig. 8e**). To assess potential regulatory roles in development, we repurposed two genetic assays previously used to characterize the START domains of PHB and CNA [28,42]. The first involves complementation of a *phb phv cna* loss-of-function mutant with YFP-tagged *pPHB:PHB* reporter constructs. Tryptophan residues in the dSTART domain of *pPHB:PHB* were replaced with alanine residues to create *pPHB:PHB-dWmut*. As expected, *phb phv cna* mutants transformed with *pPHB:PHB* were indistinguishable from wildtype plants (**Fig. 3a**). By contrast, the *pPHB:PHB-dWmut* construct failed to complement the *phb phv cna* mutant phenotype (**Fig. 3a**). This suggests PHB activity is either reduced or lost upon mutation of these tryptophan residues. To distinguish between these possibilities, we used the second genetic assay which involves transforming wildtype plants with miR166-resistant YFP-tagged *pPHB:PHB\** reporter constructs. This reporter conditions strong, conspicuous, gain-of-function phenotypes, and provides a more sensitive readout of PHB activity than complementation assays (**Supp Fig. 9**) [38,58]. Here, we similarly replaced tryptophan residues in *pPHB:PHB\** with alanine residues and transformed the resulting *pPHB:PHB*-dWmut* construct into wildtype plants. As expected, over 90% of *pPHB:PHB\** primary transformants show strong phenotypes consistent with ectopic HD-ZIPIII activity, including dorsalized leaves and enlarged meristems (**Fig. 3b**). By contrast, the majority of *pPHB:PHB*-dWmut* seedlings were indistinguishable from wildtype, with less than 30% showing weak PHB gain-of-function phenotypes (**Fig. 3b**). Thus, complementation and gain-of-function assays both indicate the dSTART conserved tryptophan residues are required for full PHB developmental function.

**Figure 3.**
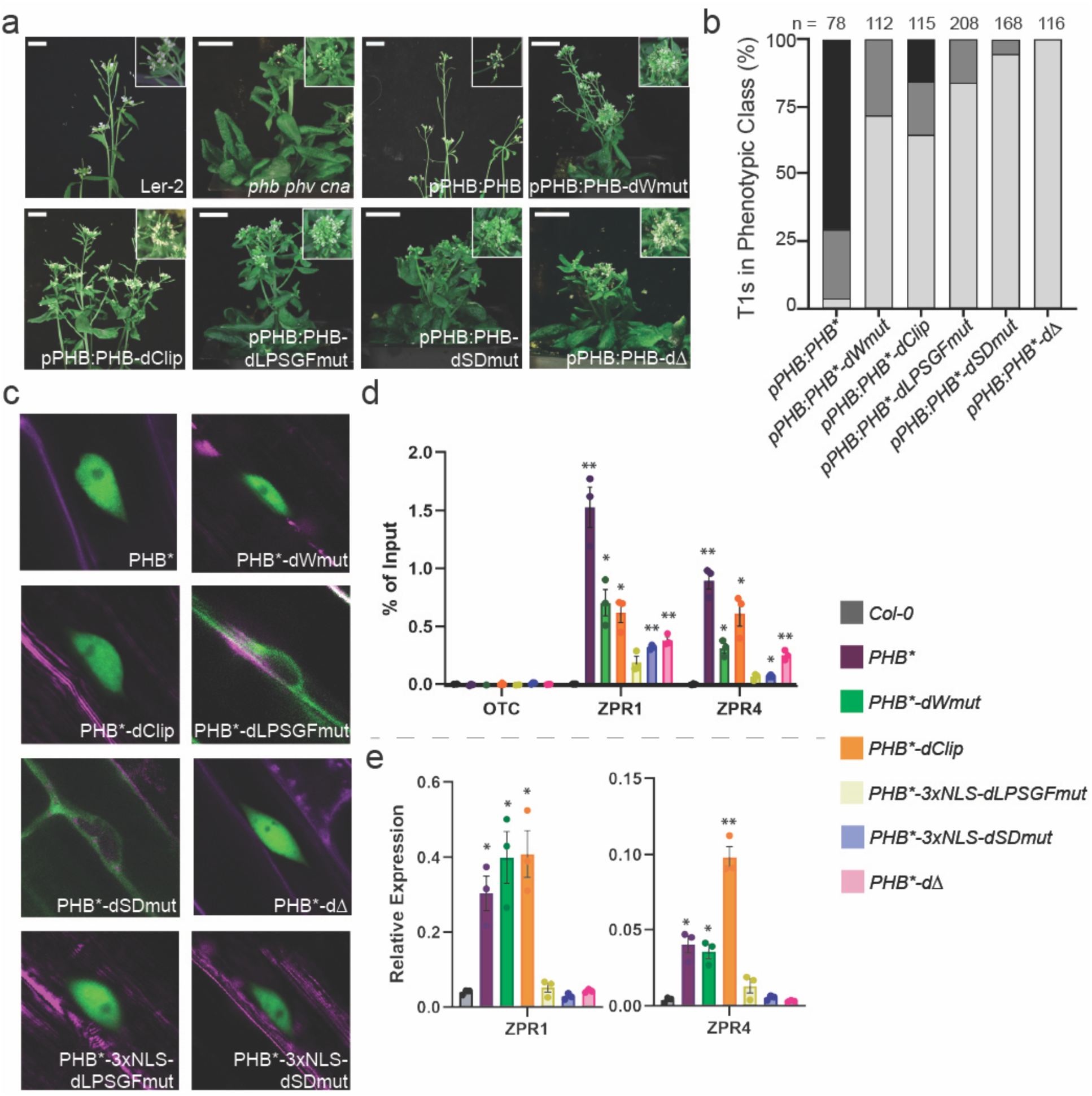
The dSTART domain controls PHB subcellular localization, DNA-binding competence, and transactivation potential. **a**, The strong pleiotropic phenotype of *phb phv cna* triple mutants is fully complemented by the *pPHB:PHB* transgene, partially complemented by the *pPHB:PHB-dClip* transgene, and unaffected by *pPHB:PHB-dWmut, pPHB:PHB-dLPSGFmut; pPHB:PHB-dSDmut*, and *pPHB:PHB-dΔ* transgenes. **b**, Phenotypic scoring of primary transformants (*n* above bar) carrying miR166-insensitive (*) constructs largely recapitulates *phb phv cna* complementation results. Categories range from severe (black) to moderate (dark grey) to mild/wildtype (light grey) phenotypes (following [28]; **Supp Fig. 9**). **c**, All PHB variants except PHB-dLPSGFmut and PHB-dSDmut are nuclear localized. Green = YFP, purple = propidium iodide. **d**, PHB binds regulatory regions of *ZPR1* and *ZPR4* and its ChIP signal is significantly enriched over the *ORNITHINE TRANSCARBAMYLASE (OTC)* negative control locus. Binding is moderately reduced for PHB-dWmut and PHB-dClip, strongly reduced for PHB-dSDmut and PHB-dΔ, and abolished for PHB-dLPSGFmut. **e**, *ZPR1* and *ZPR4* are robustly transactivated by PHB, PHB-dWmut, and PHB-dClip in 24-hour estradiol-induced lines. By contrast, PHB-dLPSGFmut, PHB-dSDmut, and PHB-dΔ fail to upregulate either target. *n* = 3 biological replicates. * = p-value ≤ 0.01, ** = p-value ≤ 0.001, two-way paired Student’s t-test.

We next cloned the *PHB*-dWmut* coding sequence into an established short-term estradiol induction system to test whether PHB-dWmut shows defects in any known StARkin-directed regulatory mechanisms [28,42]. Confocal imaging and western blotting of 24-hour estradiol-induced lines indicate PHB and PHB-dWmut accumulate in the nucleus, and to comparable levels (**Fig. 3c; Supp Fig. 10**). Thus, dSTART tryptophan residues do not control PHB activity through subcellular localization or protein stability. We then tested DNA-binding competence by assaying PHB-dWmut occupancy at established HD-ZIPIII targets using chromatin immunoprecipitation (ChIP; [28–31]). PHB-dWmut ChIP signal intensities at *ZPR1* and *ZPR4* introns were approximately half those of PHB suggesting DNA-binding competency is moderately reduced in this variant (**Fig. 3d**). We then tested whether target activation was similarly reduced. Interestingly, *ZPR1* and *ZPR4* targets were induced to comparable levels as PHB, despite reduced occupancy of PHB-dWmut (**Fig. 3e**). One possibility is that transactivation potential is moderately increased by mutation of the dSTART conserved tryptophan residues. A more likely explanation is that the elevated expression of PHB-dWmut in the estradiol-inducible lines is sufficient to overcome the effects of its hypomorphic binding capabilities. These data indicate that the tryptophan residues common to nearly all StARkin superfamily members are required for wildtype PHB DNA-binding activity and thus full PHB developmental function.

### The dSTART α3-β2 IDR-containing insertion is required for full PHB function

The dSTART domain is defined by unique structural features including an IDR-containing insertion between its α3 and β2 secondary structure elements. To test whether this 41-amino-acid-long feature contributes to PHB dSTART regulation, we clipped 36 amino acids out of its center, creating an α3-β2 linker matching those of canonical START domains (dClip; [48]). This truncation minimally affects protein folding at other regions of the dSTART domain (**Supp Fig. 8b**), and circular dichroism confirmed wildtype and dClip dSTART domains purified from *E. coli* have similar secondary structures (**Supp Fig. 8e**). Amino acids were clipped from both *pPHB:PHB* and *pPHB:PHB\** reporters to create *pPHB:PHB-dClip* and *pPHB:PHB*-dClip*, respectively. In contrast to *pPHB:PHB*, the *pPHB:PHB-dClip* transgene only partially complemented the *phb phv cna* triple mutant phenotype (**Fig. 3a**). Further, severe gain-of-function phenotypes were observed much less frequently in *pPHB:PHB*-dClip* primary transformants compared to those transformed with the *pPHB:PHB\** transgene (**Fig. 3b**). Thus, the α3-β2 insertion is required for full dSTART activity, but its effects appear to be weaker than those of the conserved dSTART tryptophan residues. The *PHB*-dClip* cassette was then cloned into the estradiol-induction system and tested for defects in StARkin regulatory mechanisms. Subcellular localization and protein stability of PHB-dClip were indistinguishable from PHB, ruling them out as regulatory mechanisms (**Fig. 3c**). However, like PHB-dWmut, PHB-dClip showed moderately reduced ChIP signal at *ZPR1* and *ZPR4* loci (**Fig. 3d**), but normal (or possibly elevated) transactivation of these targets (**Fig. 3e**). Thus, like the conserved dSTART tryptophan residues, the dSTART α3-β2 IDR-containing insertion is required for wildtype PHB DNA-binding activity and thus full PHB developmental function.

### Motifs within the dSTART domain control PHB subcellular localization

The second defining feature of the dSTART domain is the highly conserved LPSGF motif lining the interior of its hydrophobic binding pocket (**Fig. 1d**). As the SGF residues within this motif show the strongest conservation (**Supp Doc 1**), we tested their contribution to dSTART activity by mutating them into the physico-chemically similar amino acids TAY (dLPSGFmut; [59]). After confirming protein folding and secondary structure are minimally affected in dLPSGFmut (**Supp Figs 8c, 8e**), these mutations were introduced into both *pPHB:PHB* and *pPHB:PHB\** reporters to create *pPHB:PHB-dLPSGFmut* and *pPHB:PHB*-dLPSGFmut*, respectively. Interestingly, neither construct had any effect on development, with *pPHB:PHB-dLPSGFmut* failing to complement the *phb phv cna* triple mutant (**Fig. 3a**), and *pPHB:PHB*-dLPSGFmut* failing to condition any gain-of-function phenotypes (**Fig. 3b**), suggesting SGF-to-TAY mutations severely affect PHB function.

We next tested whether canonical StARkin-directed regulatory mechanisms were affected by these mutations. PHB-dLPSGFmut showed no obvious defects in protein stability (**Supp Fig. 10**), but, remarkably, appears to be completely retained in the cytoplasm (**Fig. 3c**). Multiple scenarios may explain this result. For instance, LPSGF may be a nuclear localization signal (NLS) or may mediate the response to an NLS located elsewhere on the domain. Alternatively, these residues may be involved in cytoplasmic retention. For instance, if LPSGF residues drove the uncoupling of PHB from cytoplasmic retention factors, their mutation could lead to a dominant negative protein unable to enter the nucleus.

To distinguish between these possibilities, we ran the dSTART sequence through various localization-prediction tools (**Supp Fig. 11**; see **Methods**). No NLS sequences were predicted in the dSTART domain arguing against SGF mutations directly or indirectly interfering with nuclear import. Cytoplasmic retention sequences are variable and context-dependent in nature. As such, they are not captured by traditional localization-prediction tools. However, these motifs often cluster with nuclear export signal (NES) sequences [60–62] or even act as NES motifs, themselves [63]. Strikingly, multiple strong NES sequences were predicted within the dSTART domain, including LPSGF itself (**Supp Fig. 11**). At first glance, this observation seems counterintuitive as mutating a cytoplasmic retention motif should, in principle, lead to nuclear localization. However, this can be reconciled by the presence of other predicted NES and/or cytoplasmic retention sequences within the dSTART domain, including DAPLL and RDMYLLQ. These two motifs are noteworthy because they are analogous to the amino acid stretches previously shown to control ligand-binding and DNA-binding competence in the PHB START domain [28]. Together, these findings point to a model in which cytoplasmic retention – and its subsequent relief upon ligand binding – are governed by three conserved motifs within the dSTART domain.

This hypothesis generates two testable predictions. First, mutation of DAPLL and RDMYLLQ should phenocopy PHB-dLPSGFmut, as this variant possesses a functional retention motif (LPSGF) and would be unable to relieve this cytoplasmic retention. Second, complete removal of the dSTART domain – and thus all three cytoplasmic retention sequences – should restore nuclear localization. To test the first prediction, we mutagenized DAPLL and RDMYLLQ into AVVAA and GAVVGAG, respectively (dSDmut), following previous work [28]. After confirming the integrity of dSDmut folding and secondary structure (**Supp Figs 8d, 8e**), these mutations were introduced into both *pPHB:PHB* and *pPHB:PHB\** reporters, to create *pPHB:PHB-dSDmut* and *pPHB:PHB*-dSDmut*, respectively. Neither construct affected development (**Fig. 3a**, **3b**), and, consistent with our hypothesis, PHB-dSDmut is completely retained in the cytoplasm, mirroring the accumulation pattern of PHB-dLPSGFmut (**Fig. 3c**). To test the second prediction, we removed the dSTART domain from both *pPHB:PHB* and *pPHB:PHB\** reporters, to create *pPHB:PHB-dΔ* and *pPHB:PHB*-dΔ*, respectively. Neither construct affected development (**Fig. 3a**, **3b**), suggesting compromised function of PHB-d*Δ*. Most importantly, the accumulation pattern of PHB-d*Δ* is strictly nuclear, consistent with the loss of cytoplasmic retention sequences (**Fig. 3c**). Taken together, these data suggest three cytoplasmic retention sequences within the dSTART domain – LPSGF, DAPLL, and RDMYLLQ – regulate HD-ZIPIII subcellular localization, possibly in response to binding of a ligand.

### Motifs within the dSTART domain control PHB DNA-binding competence

StARkin domains impact biological processes through a diverse set of regulatory mechanisms, and a given StARkin domain often uses multiple independent mechanisms (reviewed in [37]). For instance, the START domain of PHB impacts homodimerization, transactivation potential, and DNA-binding competence [28]. It is thus conceivable that some of these properties are also affected by the dSTART domain. Alternatively, dSTART activity may be limited solely to control of subcellular localization. If so, PHB-dLPSGFmut and PHB-dSDmut would be fully functional proteins, apart from their cytoplasmic retention phenotype. To distinguish between these possibilities, we fused a *3x-NLS* tag to the N-termini of *PHB-dLPSGFmut* and *PHB-dSDmut* cassettes to override their cytoplasmic retention, allowing us to assess possible effects on transcriptional activity. Cassettes were cloned into the estradiol-inducible system, and confocal imaging confirmed that both 3xNLS-PHB-dLPSGFmut and 3x-NLS-PHB-dSDmut accumulate in the nucleus, as expected (**Fig. 3c**). We then tested the DNA-binding capabilities of these variants by measuring occupancy at *ZPR1* and *ZPR4* loci. Strikingly, ChIP signal intensities were sharply reduced for 3xNLS-PHB-dLPSGFmut and 3x-NLS-PHB-dSDmut relative to PHB, particularly at the *ZPR4* locus (**Fig. 3d**). In addition, both variants fail to upregulate *ZPR1* and *ZPR4* targets suggesting this reduced binding capacity has functional consequences or that these mutations also reduce transactivation potential (**Fig. 3e**). Taken together, our findings propose the LPSGF, DAPLL, and RDMYLLQ motifs within the dSTART domain contribute to HD-ZIPIII DNA-binding competence and target regulation, in addition to their control of HD-ZIPIII subcellular localization.

### The dSTART domain facilitates the differential usage of shared binding sites

Functional divergence of two HD-ZIPIII paralogs – CNA and PHB – results from differential usage of shared binding sites, with hundreds of uniquely regulated targets emerging from a commonly bound genetic network [42, reviewed in 64]. Deletions, chimeras, and evolutionarily-guided amino acid substitutions show that START domains partially license this phenomenon by helping to distinguish responsive from non-responsive binding sites [42]. Given their evolutionary relatedness (**Fig. 2**), it is conceivable the dSTART domain also contributes to the differential usage of shared binding sites by HD-ZIPIII TFs.

To address this, we used 24h-estradiol-induced lines to test how loss of the dSTART domain affects binding and regulation of PHB targets genome wide. ChIP-seq identified 3049 and 2999 sites bound by PHB and PHB-dΔ, corresponding to 3093 and 3044 bound genes, respectively (**Figs 4a, 4b, 4c**). The binding profiles of PHB and PHB-dΔ are virtually identical, with ∼98% of genes bound by PHB also bound by PHB-dΔ (3044 out of 3093). However, clear differences were observed in ChIP signal intensities across their 2999 mutually bound sites. ChIP signal intensities fell into three categories: higher signal for PHB (1863 out of 2999), higher signal for PHB-dΔ (1 out of 2999), and equivalent signal for both variants (1135 out of 2999; **Figs 4a, 4b**). Thus, like the PHB and CNA START domains [42], one function of the dSTART domain may be to increase DNA binding affinities in a site-specific manner.

**Figure 4.**
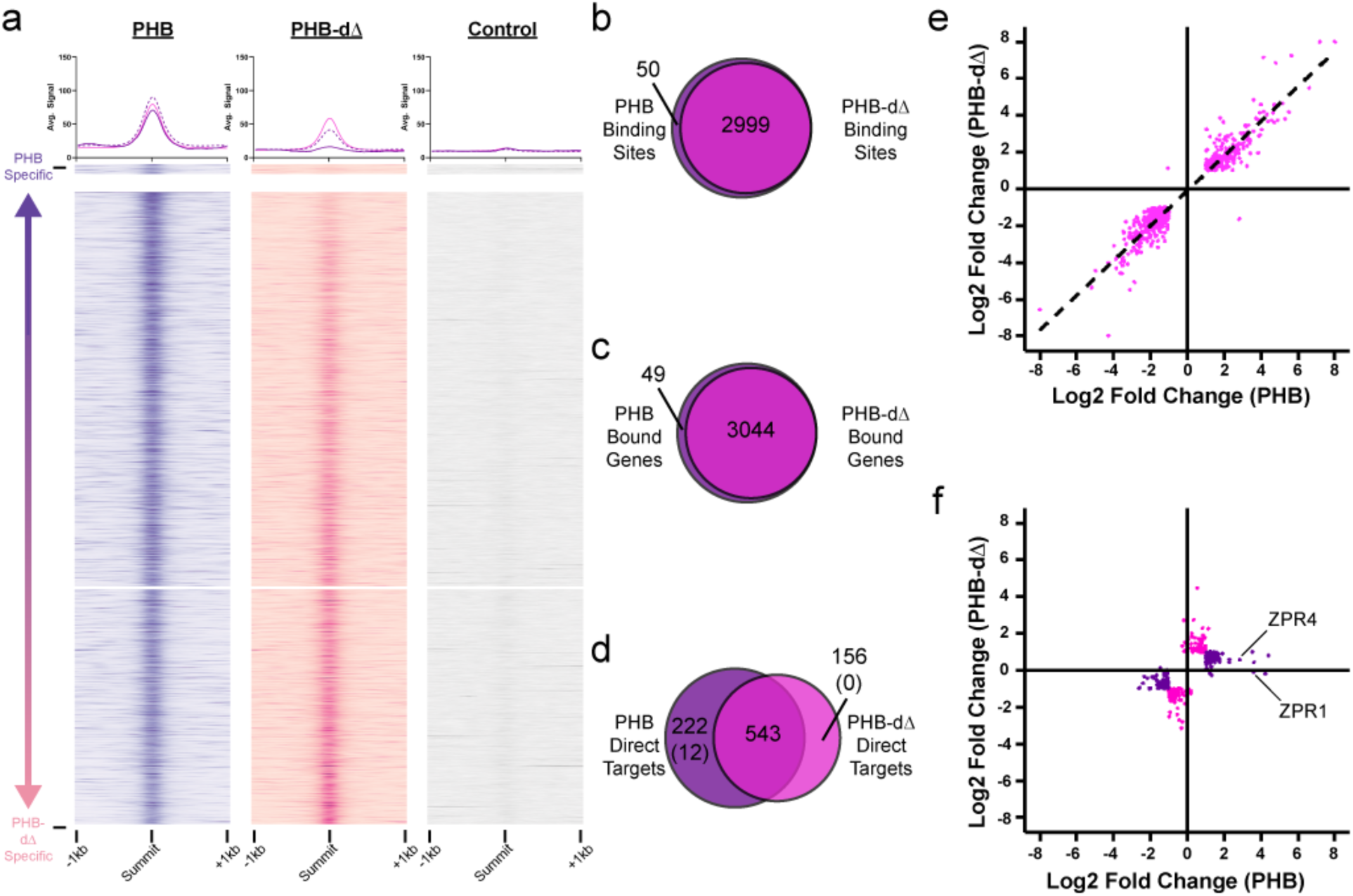
The dSTART domain helps distinguish responsive from non-responsive PHB binding sites. **a**, Histograms and heatmaps of ChIP-seq signal intensities compared to a non-transgenic control (grey). Histogram lines and colors delineate four categories of binding sites: PHB uniquely bound (purple, solid); mutually bound – PHB higher affinity (purple, dashed); mutually bound – equal affinities (magenta); and mutually bound – PHB-dΔ higher affinity (pink, dashed). Heatmaps are separated into two major categories: PHB uniquely bound (upper) and mutually bound (lower). Mutually bound heatmaps contain two further subcategories: mutually bound – PHB higher affinity (upper); and mutually bound – equal affinities (lower). **b**, Venn diagram of sites bound by PHB and PHB-dΔ. **c,** Venn diagram of genes bound by PHB and PHB-dΔ. **d,** Venn diagram of direct targets of PHB and PHB-dΔ, i.e. bound in ChIP-seq and differentially expressed in RNA-seq. Numbers in parentheses correspond to direct targets bound specifically by PHB and PHB-dΔ. **e**, Scatterplot showing differential expression of genes bound and regulated by both PHB and PHB-dΔ. **f**, Scatterplot showing differential expression of genes bound by both paralogs but uniquely regulated by PHB (purple) or PHB-dΔ (pink).

We then tested how loss of the dSTART domain impacts regulation of PHB targets using RNA-seq. After induction, 1758 and 1707 genes were differentially expressed in PHB and PHB-dΔ lines, respectively (**Supp Tables 2b, 2c**). Genes that were also bound in ChIP assays were called as direct targets. This corresponds to 777 and 699 high-confidence direct targets of PHB and PHB-dΔ, respectively, which fall into three categories: PHB-specific, PHB-dΔ-specific, and mutually regulated (**Fig. 4d**). Several mechanistic conclusions can be drawn from these data. First, mutual targets of PHB and PHB-dΔ are almost all regulated in the same direction (**Fig. 4e**). This indicates the dSTART domain does not control the valence of PHB target regulation. Second, the magnitudes of gene expression changes induced by PHB and PHB-dΔ are similar, demonstrating that deletion of the dSTART domain does not impair transcriptional potency. This stands in contrast to the START domain whose loss reduces transcriptional potency of both CNA and PHB [28,42]. Related to this, the majority of PHB and PHB-dΔ direct targets fall into the mutually regulated category, despite the reduced binding affinity of PHB-dΔ (**Fig. 4d**). This indicates that, like PHB-dWmut and PHB-dClip (**Fig. 3d**), the effects of PHB-dΔ hypomorphic DNA binding can be overcome by increasing dosage. Third, despite having virtually identical binding profiles, PHB and PHB-dΔ each have hundreds of uniquely regulated targets (**Fig. 4f**). Specifically, 27% of PHB-specific targets (210 out of 765) and 22% of PHB-dΔ-specific targets (156 out of 699) are also bound by the other variant. These PHB-specific targets include *ZPR1* and *ZPR4*, which remain bound by PHB-dΔ, but are not upregulated by this variant (**Fig. 4f**). As transcriptional potency is not reduced at all PHB-dΔ targets (**Fig. 4e**), this suggests the dSTART domain functions by helping to discriminate between responsive and non-responsive binding sites. Taken together, these and previous findings [42] demonstrate the differential usage of shared binding sites by HD-ZIPIII TFs is mediated by both their START and dSTART domains.

### The dSTART domain binds phosphatidylglycerol and phosphatidic acid

A core property of StARkin domains is their binding to hydrophobic ligands [34–36], which can have important regulatory consequences [reviewed in 37]. For instance, the PHB START domain binds to multiple species of PC, and binding of PC is required for PHB DNA-binding competence [28]. This prompted us to test whether the dSTART domain similarly binds to lipophilic ligands. To do this, we used a tandem-purification strategy to obtain highly-pure GFP-tagged recombinant dSTART protein from *E. coli* (**Supp Fig. 12a**; see **Methods**). GFP-tagged PC-transfer protein (PC-TP) and GFP alone were purified in parallel as controls (**Supp Fig. 12a**). Three lines of evidence support functionality of the recombinant proteins. First, isolated proteins were strongly fluorescent, consistent with a properly folded GFP tag. Second, circular dichroism showed dSTART and PC-transfer protein (PC-TP) have similar secondary structures, which are distinct from the GFP negative control (**Supp Fig. 8f**). Third, liquid-chromatography mass spectrometry (LC-MS) confirmed binding of expected bacterial lipids. Here, lipids were extracted from GFP and dSTART proteins, subjected to LC-MS, and queried against a panel of over 1,200 species of phospholipids (**Supp Table 3**). Multiple phosphatidylglycerol (PG) species were highly enriched across the four dSTART replicates (≥ 3-fold enrichment, *q* ≤ .01; **Fig. 5a**), and these species match previously reported “fortuitous bacterial ligands” bound by other recombinant phospholipid binding proteins, including PC-TP [65], steroidogenic factor 1 (SF-1, [66]), liver receptor homolog-1 (LRH-1, [67]), and the HD-ZIPIII START domain [28].

**Figure 5.**
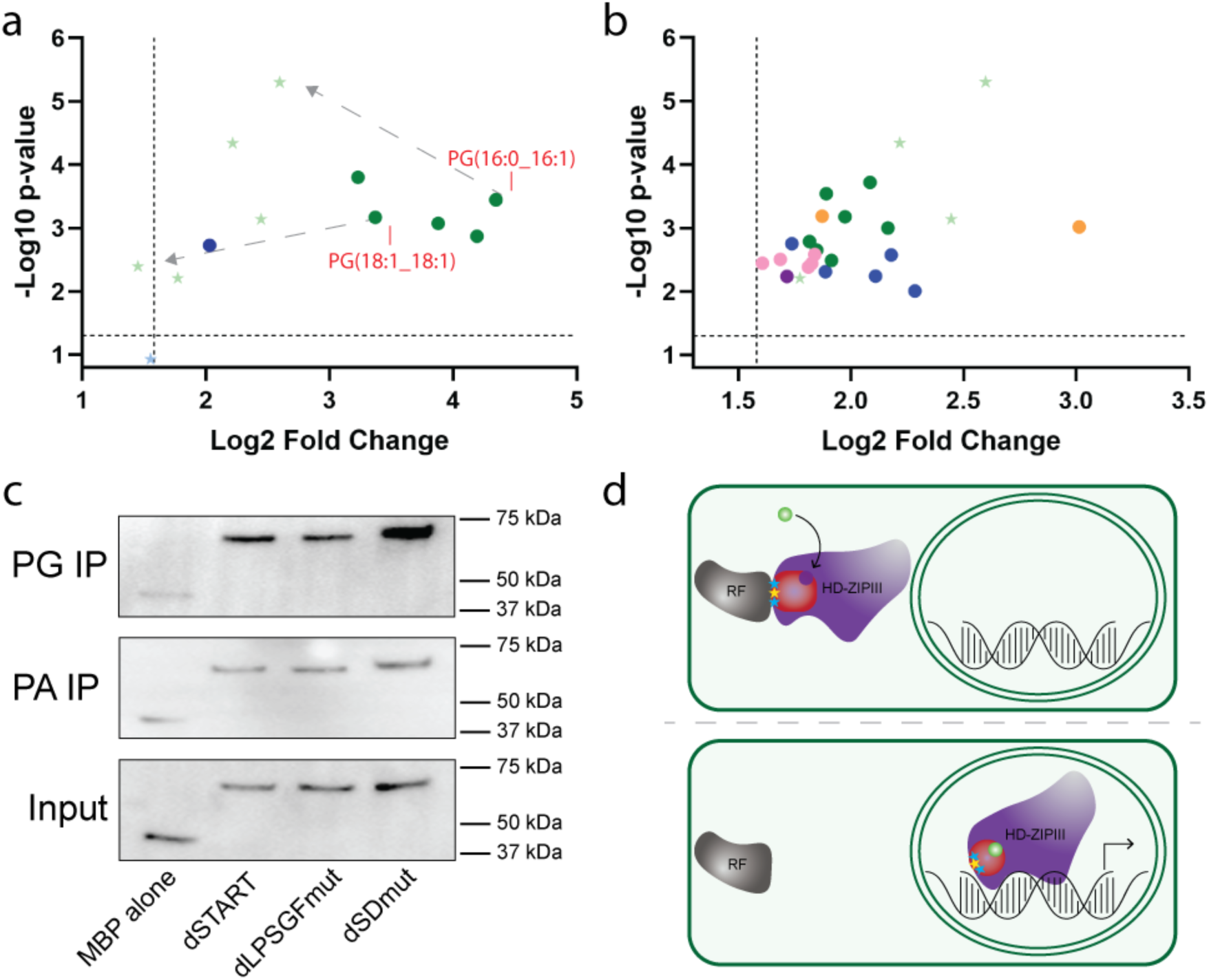
The dSTART domain binds multiple species of phosphatidylglycerol (PG) and phosphatidic acid (PA). **a**, LC-MS analysis of lipids bound by recombinant dSTART protein in bacteria. Five species of PG (green circles) and one of PA (blue circle) are significantly enriched (≥ 3-fold enrichment, *q* ≤ .01) over the negative control and are depleted after incubation with plant liposomes (light green or blue stars). Two representative lipids, before and after liposome incubation, are labeled with red text and grey arrows. **b**, Twenty new lipid species are significantly enriched (≥ 3-fold enrichment, *q* ≤ .01) after incubation with plant liposomes including: PG (green circles), PA (blue circles), cardiolipins (CL; pink circles), PC (orange circles), and phosphatidylinositol (PI; purple circle). Four species of PG also bound in bacteria remain significantly enriched after liposome incubation (green stars). **c**, dSTART, dLPSGFmut, and dSDmut proteins bind to PG- and PA-conjugated beads. Binding to PG is noticeably stronger than binding to PA. **d**, Working model of dSTART regulation of HD-ZIPIII activity. Retention factors (RF; grey) hold HD-ZIPIII TFs (purple) outside of the nucleus via recognition of LPSGF (yellow star) or SDmut (DAPLL, RDMYLLQ; blue stars) sequences. Nuclear exclusion is relieved by conformational change of the dSTART domain triggered by binding of its PG or PA ligands (green circle). The dSTART domain also promotes DNA-binding competence or affinity and contributes to the differential usage of shared binding sites by HD-ZIPIII TFs through an unknown mechanism.

After confirming functionality, GFP and dSTART proteins were incubated with liposomes generated from *Arabidopsis* total lipid extracts, repurified by affinity chromatography, and subjected to LC-MS. Consistent with efficient exchange for endogenous plant ligands, most bacterial ligands were strongly depleted after liposome incubation (**Fig. 5a**), and twenty new lipids were enriched specifically in dSTART samples (≥ 3-fold enrichment, *q* ≤ .01; **Fig. 5b**). Seven of these new lipids are PGs and five are members of the phosphatic acid (PA) class. Also enriched were five species of cardiolipin (CL) – phospholipid dimers comprised of monomers of PG and PA [68]. Thus, out of the twenty identified plant-enriched lipids, seventeen of them have either PG or PA as part of their structure (**Fig. 5b**). To independently confirm binding, we incubated recombinant MBP-tagged dSTART protein with PG- or PA-conjugated beads. dSTART bound both phospholipid classes (**Fig. 5c**), and binding was stronger to PG, mirroring its greater recovery in LC-MS analyses. Finally, we tested the impact of LPSGF, DAPLL, and RDMYLLQ motifs on PG and PA binding using recombinant dLPSGFmut and dSDmut proteins. Interestingly, both proteins retain PG- and PA-binding capacity (**Fig. 5c**). These residues may thus mediate the conformational response induced by ligand binding, rather than binding itself, a phenomenon that has been observed for other StARkin domains [34]. A non-mutually exclusive explanation is that these residues do directly participate in ligand binding but redundantly with many others, as seen for StARkin domains such as PC-TP [35]. Taken together, our analyses propose PG and PA phospholipids as high-confidence candidate ligands of the dSTART domain.

## DISCUSSION

The StARkin domain is a ubiquitously distributed evolutionary module that uses a diverse set of regulatory mechanisms to impact a wide range of biological processes [37]. Here we demonstrate that HD-ZIPIII and HD-ZIPIV TFs contain a cryptic StARkin domain defined by multiple unique features including a highly conserved LPSGF amino acid motif and an IDR insertion between its α3 and β2 secondary structure elements. This new domain, which we have termed the disorder-containing START (dSTART) domain, is present in the earliest known HD-ZIPIII and HD-ZIPIV TFs, indicating its origin predates the divergence of streptophyte algae over 725 million years ago [69,70]. Using PHB as a representative member, we show the dSTART domain is required for full HD-ZIPIII developmental function. Evolutionarily-guided mutagenesis identified multiple distinct dSTART-regulated mechanisms, including promotive effects on PHB nuclear localization and DNA binding competence and/or affinity. Genomic analyses indicate the dSTART domain also contributes to the differential usage of shared binding sites phenomenon previously observed for PHB and CNA [42]. Finally, we identify members of the PG and PA phospholipid classes as high-confidence candidate ligands of the PHB dSTART domain. Our findings propose a model in which phospholipid binding at the HD-ZIPIII dSTART domain relieves nuclear exclusion, promotes occupancy across a shared network architecture, and helps to generate paralog-specific transcriptional outcomes (**Fig. 5d**).

Some regulatory mechanisms are shared by the START and dSTART domains. For instance, both StARkin domains show promotive effects on DNA-binding competence or affinity. Here, mutations preventing PC-binding by the START domain abolish binding of PHB to *ZPR3* and *ZPR4* loci [28], while 3xNLS-PHB-dLPSGFmut and 3x-NLS-PHB-dSDmut show sharply reduced binding at their *ZPR1* and *ZPR4* targets. Similarly, deletion of either StARkin domain reduces ChIP signal across hundreds of loci, an effect which is stronger for the dSTART domain (**Fig. 5**; [42]). START and dSTART domains also play roles in mediating the differential usage of shared binding sites phenomenon, as loss of either domain drives the interconversion of hundreds of responsive and non-responsive binding sites[42]. Logical next steps include determining whether each StARkin domain licenses its own unique subset of shared sites as well as testing for genes whose expression depends on allostery between the domains. The underlying drivers of this understudied mechanism of TF functional divergence remain completely unknown in plants and animals. However, to date, all TFs shown to use this mechanism contain ligand binding domains [42,71]. It is therefore tempting to speculate that ligands may be involved in mediating binding site discrimination. An alternate, though not mutually exclusive mechanism could be protein partners whose interaction is directly or indirectly controlled by the StARkin domains. Direct interactions between proteins and StARkin domains are well-documented [72–80], making the dSTART α3-β2 insertion – a potential IDR-to-α-helix switch – particularly intriguing.

Other regulatory mechanisms used by the START and dSTART domains are distinct. For instance, the START domain increases PHB transcriptional potency, which does not seem to be the case for the dSTART domain [28]. Further, mutation, deletion, and sequence analyses show no evidence of START-dependent effects on subcellular localization (**Supp Fig. 11**; [28]). By contrast, the dSTART domain contains three predicted NES and/or cytoplasmic retention sequences including the highly conserved LPSGF motif that defines this new subfamily. Our findings are consistent with a model in which the *apo* form of the dSTART domain holds PHB outside of the nucleus, perhaps through interaction with cytoplasmic, cytoskeletal, or membrane components. Conformational response driven by binding of the dSTART ligand would relieve this inhibition allowing PHB to translocate to the nucleus. Multiple metazoan NRs serve as precedence for this regulatory paradigm, including the mineralocorticoid receptor (MR), the androgen receptor (AR), and the estrogen receptor α (ERα), whose NES sequences contain ligand-contacting residues and whose nuclear import is triggered by ligand binding [81–83].

In addition to their distinct sets of regulatory mechanisms, the START and dSTART domains also have distinct lipid binding profiles. For instance, unlike the PHB START domain, no species of phosphatidylethanolamine (PE) were significantly enriched in bacterial or plant dSTART samples (**Supp Table 3**; [28]). Further, only two species of PC were recovered after incubation of dSTART with plant liposomes (**Fig. 5b**), and neither match previously identified START ligands [28]. Finally, out of the twenty plant-enriched dSTART lipids, only PG(34:1) was also significantly enriched in START samples [28]. Thus, the START and dSTART domains both bind to phospholipid ligands but appear to have markedly different class and species preferences.

A remarkable 85% of the dSTART ligands recovered after incubation with *Arabidopsis* liposomes contained either PG or PA. Links between PA- or PG-binding and TF subcellular localization have been noted in other systems, including plants. For instance, binding of PA triggers nuclear translocation of the R2R3 MYB TF WEREWOLF (WER; [84]). Mutations that prevent PA binding led to cytoplasmic accumulation of WER and the specification of hair cells in place of nonhair cells. Interestingly, when driven to the nucleus by the addition of an NLS tag, these mutant WER variants were fully functional [84]. This distinguishes them from PHB-dLPSGFmut and PHB-dSDmut and highlights the fact that multiple regulatory mechanisms are controlled by the LPSGF, DAPLL, and RDMYLLQ motifs. In another example, the mobile florigen molecule FLOWERING LOCUS T (FT) binds to PG, and, at low temperatures, this binding anchors FT to intracellular membranes [85]. As temperatures increase, binding is disrupted, releasing FT from the membrane and allowing it to traffic to the shoot apical meristem to promote the floral transition. These examples illustrate phospholipids can act as environmental sensors, and that spatial regulation of their accumulation is an efficient way to pattern TF activity. In our experiments, we did not observe overt non-nuclear accumulation of wildtype PHB protein, suggesting PA and PG are not limiting under the conditions and cell types we tested. Careful examination of HD-ZIPIII subcellular localization across a wide range of developmental and environmental contexts is thus an important next step.

Artificial neural networks like AlphaFold2 are an increasingly important tool to predict function from structure. For instance, elegant work using AlphaFold2 multimeric models of full-length HD-ZIPIII and HD-ZIPIV proteins suggests different modes of action for their dSTART domains [43]. For some HD-ZIPIV proteins, homodimerization is proposed to complete the lipid binding pocket of each dSTART monomer. Binding of dSTART ligands would therefore be stabilized by an interface formed by both dSTART domains. This scenario would not have emerged from homology modeling of the dSTART domain in isolation. By contrast, HD-ZIPIII multimers are not predicted to homodimerize via either of their START domains [43]. In addition to highlighting the power of modeling to inform biological hypotheses, these findings nicely illustrate the dissimilarities between the HD-ZIPIII and HD-ZIPIV subfamilies, despite their evolutionary relatedness. For instance, HD-ZIPIV START domains can bind to sphingolipids (i.e. very long-chain fatty acid-containing ceramides (VLCFA-Cers; [86]) while HD-ZIPIII StARkin domains seem to prefer phospholipids (i.e. PC, PG, or PA; [28]). In addition, DNA-binding activity is not affected by the HD-ZIPIV START domain [39–41,87], whereas this is a key regulatory feature of both HD-ZIPIII StARkin domains [28,42]. Further, proteasome-dependent protein turnover is a major regulatory control point for HD-ZIPIV START domains, which careful experiments indicate is intricately tied to binding of their VLCFA-Cers and lyso-PC ligands [86]. By contrast, there is no evidence this regulatory mechanism is affected by either HD-ZIPIII StARkin domain. Finally, evolutionary conservation patterns also differ between the subfamilies, with the HD-ZIPIII START and HD-ZIPIV dSTART domains showing higher conservation than their adjacent StARkin domain. The significance of this is unclear but may reflect different selective pressures on distinct regulatory properties.

In summary, we report the identification and functional characterization of a cryptic, deeply conserved StARkin domain contained within key regulators of plant development. Our findings reinforce the flexible and diverse nature of the StARkin regulatory module and open new research directions. These include testing for allostery between the START and dSTART domains, identifying drivers of dSTART-mediated effects on subcellular localization and differential usage of shared binding sites, and functionally characterizing the HD-ZIPIV dSTART domain whose suite of regulatory mechanisms remain unknown.

## METHODS

### Bioinformatics tools and databases

PSI-BLAST searches against the RefSeq database at NCBI were carried out with default parameters. Protein domain architectures were obtained from the Pfam (30357350), Conserved Domain Database (25415356), and TREND (32282909) servers. Profile to profile searches for domain identification were carried out in HHpred (15980461). To collect homologous sequences, full-length sequence of *A. thaliana* PHB protein was used as a query for BLASTP searches against the genomes of the representative set of Viridiplantae species available in NCBI RefSeq and NCBI non-redundant databases. Additionally, the 1KP database was scanned for the HD-ZIPIII proteins in Chlorophyta lineage.

### Homology modeling, pairwise similarity studies, and multiple sequence alignments

Modeling of the dSTART domain was performed by I-TASSER (https://zhanggroup.org/I-TASSER) and AlphaFold2 (https://alphafold.ebi.ac.uk) using default parameters. Models were visualized in Pymol. Pairwise comparisons between different START domains done via MAFFT (Multiple Alignment using Fast Fourier Transform) and the percent identity matrix readout. Values were input into Microsoft Excel and colored through the heatmap function. For minimal red/blue multiple sequence alignments, PHB dSTART paralog amino acid sequences were aligned using MAFFT (https://mafft.cbrc.jp/alignment/server). Multiple sequence alignments were constructed using LINS-I and FFT-NS-2 algorithms in MAFFT (28969743). Sequence alignments were edited in Jalview (19151095). Secondary structure prediction from amino acid sequences were carried out using Ali2D and Quick2D from the MPI Bioinformatics Toolkit (29258817). Blue and red MSAs had conserved aligned residues with a 98% consensus threshold identified using R and the “msa” package from Bioconductor. The file was exported in .tex format and formatted via TeXworks.

### Plant materials and growth conditions

*Arabidopsis thaliana* (Col-0 ecotype) seedlings were grown on soil at 22°C in long day conditions on soil or on 1% agarose Murashige and Skoog medium plates (pH 5.7). 24-hour estradiol inductions were performed by spraying 7- or 10-day-old seedlings on plates with 50 µM B-estradiol in 1% DMSO and 0.005% Silwet.

### Molecular biology and plant transformations

The pPHB:PHB and *pPHB:PHB\** constructs have been described previously and were used as templates to create the complementation and gain-of-function/overexpression constructs, respectively. Gain-of-function constructs were created as follows. Using Gibson assembly (NEB), to create *pPHB:PHB*-dΔ*, the 867bp dSTART domain (1191-2058bp from the start codon) was removed through overlapping primers containing linker regions upstream and downstream of the dSTART domain. To create *pPHB:PHB*-dWmut*, we synthesized a mutated dSTART domain variant (GeneArt), which made amino acid changes W425A and W492A by substituting the W codon with GCG. To create the *pPHB:PHB*-dClip* construct, overlapping primers were made to remove amino acids 605 to 645 (1819-1926bp from the start codon). Similarly, the *pPHB:PHB*-dSDmut* construct was created through a synthesized mutated dSTART domain variant (Twist Bioscience), which made amino acid changes of RDMYLLQ and DAPLL to AVVAAV and AVVAA, respectively. Finally, the *pPHB:PHB*-dLPSGFmut* construct was made by substituting the SGF amino acids (positions 599-601) with TAY (ACCGCATAC). All constructs were then placed into the pB7GW binary vector via Gateway LR reactions (VIB Ghent; Invitrogen). Complementation assays were similarly constructed but using miR166-sensitive *pPHB:PHB* as a template.

For the estradiol-induction constructs, the coding sequence regions of all miR166-resistant constructs were reamplified and cloned into pCR8-GW entry vector using Gibson assembly. Gateway LR reactions were then performed to place the coding regions into the *pUBQ10:XVE oLexA* system described in [28].

All constructs, both native and inducible, were transformed into plants using *Agrobacterium tumefaciens*-mediated floral dip transformation. Cloning primers and synthesized sequences can be found in **Supp Table 4**.

### Confocal imaging and western blotting

Induced seedlings (5 days old) were incubated in 0.5 µg/mL propidium iodide solution for 10 seconds to stain the root cell plasma membranes and then washed in water to remove excess propidium iodide, as described previously [28]. Full seedlings were wet mounted to a glass slide with a coverslip. Roots were examined under a Leica Stellaris 5 using a Leica C plan 40x objective. The mCitrine YFP tag and propidium iodide were excited using the 526 nm wavelength, and emission was captured between 490 and 550 nm or 570 and 620 nm, respectively.

For western blotting, lysates from induced seedlings were separated on an SDS-PAGE gel, blotted to nitrocellulose membrane, blocked with 5% milk fat in TBST for 1 hour at room temperature, then probed with 1:1000 anti-FLAG (Millipore) or 1:1000 anti-Actin (Millipore). Proteins were visualized using 1:5000 HRP-conjugated secondary antibody (Jackson) and SuperSignal West Pico PLUS substrate (Pierce).

### Chromatin immunoprecipitation and qPCR

10-day old seedlings were grown densely on MS plates and sprayed with estradiol to induce expression. 24 hours later, seedlings were collected off the plates and placed in 1x PBS containing 2% formaldehyde (ThermoFisher). Seedlings were crosslinked in a vacuum chamber for 15 minutes at 20-25 mmHg vacuum pressure. Pressure was released, tubes containing the seedlings were shaken to redistribute both seedlings and formaldehyde, then vacuum was reapplied for a further 15 minutes. Crosslinking was quenched by adding 2 mL of 2M glycine, shaking, then vacuuming for 5 additional minutes. Seedlings were rinsed three times with water, dried with paper towels, then weighed and separated into 0.5 g aliquots for flash freezing in liquid nitrogen.

Seedlings were ground under liquid nitrogen using a ceramic mortar and pestle, and the powdered tissue was transferred to a 50 mL conical tube on ice. 8.5 mL of nuclear isolation buffer (10mM HEPES pH 8.0, 1M sucrose, 5mM KCl, 5mM MgCl_2_, 0.6% Triton, 0.4mM PMSF, 1x cOmplete^TM^EDTA free protease inhibitor) was added to the powder and the tubes were rotated for 15 minutes at 4°C. The homogenized solution was then filtered through two layers of Miracloth into a fresh conical tube and centrifuged at 3000xg for 15 minutes at 4°C. Nuclear pellets were resuspended in 1 mL of nuclear wash buffer (10mM Tris pH 8.0, 250mM sucrose, 10mM MgCl_2_, 1mM EDTA, 1% Triton, 1mM PMSF, 1x cOmplete^TM^ EDTA free protease inhibitor) and transferred to a 1.5 mL Eppendorf tube. Tubes were centrifuged for 10 minutes at 12,000xg at 4°C and the supernatant was discarded. Pellets were resuspended in 1 mL of nuclear lysis buffer (20mM Tris pH 8.0, 2mM EDTA, 0.01% SDS, 1mM PMSF, 1x cOmplete^TM^ EDTA free protease inhibitor) and transferred to a 1 mL Covaris milliTUBE with AFA Fiber. Samples were sonicated in a Covaris S220 (150W peak power, 20% duty factor, 200 cycles/burst for 6 minutes). The sonicated chromatin was transferred to a fresh 1.5 mL Eppendorf tube and centrifuged for 10 minutes at 12,000 RPM at 4°C to pellet the insoluble debris. Soluble chromatin fractions (920 µL) were moved to a fresh 1.5 mL Eppendorf tube and 30 µL of 5M NaCl and 20 µL of 30% Triton were added to the fractions. 420 µL of sonicated chromatin was added to each IP tube and 42 µL of sonicated chromatin was put aside at -20°C for a 10% input control. 2 µL of either M2 mouse monoclonal anti-FLAG (ThermoFisher) or mouse IgG (Cell Signaling Technologies) negative control antibodies were added to their respective tubes and rotated slowly overnight at 4°C.

40 µL of Protein A/G magnetic beads (Invitrogen) were equilibrated with the nuclear lysis buffer and then added to each IP tube and rotated slowly for 2 hours at 4°C. Beads were then washed twice, with 5-minute rotations for each wash, at 4°C. The wash buffers are as follows, in order of wash: Low Salt Wash Buffer (150mM NaCl, 0.1% SDS, 1% Triton, 2mM EDTA, 20mM Tris pH 8.0), High Salt Wash Buffer (500mM NaCl, 0.1% SDS, 1% Triton, 2mM EDTA, 20mM Tris pH 8.0), LiCl Wash Buffer (250mM LiCl, 1% IGEPAL, 0.1% SDS, 1mM EDTA, 10mM Tris pH 8.0), and TE Buffer (10mM Tris pH 8.0, 1mM EDTA, 0.1% IGEPAL). Bound protein was eluted from beads by rotating with 250 µL of elution buffer (1% SDS, 0.1M NaHCO_3_) at 65°C for 15 minutes. The eluted protein-DNA complexes were removed from the beads and added to a new tube containing 10 µL of 5M NaCl. 250 µL of Elution Buffer and 10 µL of 5M NaCl was also added to the input control tubes. All tubes were placed at 65°C overnight to reverse the crosslinking. The next day, 10 µL of 0.5M EDTA, 20 µL of 1M Tris pH 7.0, 1 µL of 20 mg/mL Proteinase K, and 1 µL Rnase (50 ng/µL) were added to each tube and incubated at 45°C for 1 hour. Samples were then cleaned up with the Zymo DNA Clean and Concentrator-25 kit and eluted in 100 µL of the kit’s elution buffer. Recovered DNA was quantified using qPCR with 10 µL SsoAdvanced Universal SYBR Green Supermix (Bio-Rad), 1 µL of combined 10 µM primers, 5 µL dH_2_O, and 4 µL of DNA with two technical replicates for each tested target (OTC, ZPR1, ZPR4). Data was plotted using Prism graphing software (GraphPad).

### ChIP-seq and bioinformatics

ChIPseq libraries were constructed using the NEBNext Ultra II DNA library kit and single-end sequenced using the Illumina NextSeq1000 located on-site at the Penn Epigenetics Institute. All analyses were performed as described in [42], however key parameters include: peak calling using a combination of Model-based Analysis of ChIP-Seq (MACS2: *q* < .001) [88,89] and Irreproducibility Discovery Rate (IDR threshold: < .01) [90], and quantification of ChIP signal intensities using Diffbind (*q* < 0.05) [91]. Diffbind outputs and mapping statistics can be found in **Supp Table 2a, 2d**.

### RNA-seq and bioinformatics

Total RNA from 24-hour induced seedlings was isolated via the Direct-zol RNA Miniprep kit (Zymo). Libraries were then made via the QuantSeq 3’ mRNA-Seq V2 Library Prep Kit FWD with UDI (Lexogen) and PCR Add-on and Reamplification V2 kits (Lexogen). Libraries were then sequenced using the Illumina NextSeq1000 located on-site at the Penn Epigenetics Institute. All analyses were performed as described in [42], however key parameters include: carefully timed inductions to minimize circadian effects, application of the lfcShrink function using “ashr” [92], and calling of differentially expressed genes via DESeq2 (≥ 2 or≤ -2fold change in levels, and a *q*-value ≤ 0.1). DEseq2 comparisons and mapping statistics can be found in **Supp Table 2b, 2c, 2e**.

### qRT-PCR assays

Total RNA was extracted from 7-day-old induced *Arabidopsis* seedlings via RNAzol RT (Sigma-Aldrich) extraction methods. RNA was quantified via nanodrop and converted into cDNA via SuperScriptIV Reverse Transcriptase (ThermoFisher), using 500ng of total RNA and oligo(dT)_18_ primer. The cDNA was diluted 1:1 with Ultra-Pure H_2_O (Invitrogen) before use in qRT-PCR reactions. At least three biological replicates and two technical replicates were used for relative quantification. ΔCt values were normalized to ACTIN2 and a Student’s t test was used to calculate statistical significance.

### Protein purification, circular dichroism, and PG/PA-conjugated bead assays

Wild type dSTART (1191-2058 bp from the start codon) domain or mutant versions of the dSTART domain were cloned downstream of mEGFP in the pASG vector backbone (ThermoFisher) or downstream of MBP in a modified pET28b vector [93] via Gibson cloning. BL21 bacteria were transformed and grown to an OD of ∼0.4 to 0.6 at 37°C and induced overnight at 16°C with 100 ng/mL anhydrotetracycline for mEGFP-tagged proteins or with 100µM IPTG for MBP-tagged proteins. For mEGFP-tagged proteins, bacteria were lysed via sonication (100 mM Tris, pH 8.0, 200 mM NaCl, 2x HALT protease inhibitor) and purified on a Ni^2+^ column (Qiagen). Proteins were eluted using 250 mM imidazole diluted in lysis buffer. This protein was then doubly purified by running it over a StrepTactin Sepharose (IBA) column, washing, and eluting using 2.5 mM desthiobiotin diluted in lysis buffer. For MBP-tagged proteins, bacteria were lysed (100 mM Tris, pH 8.0, 200 mM NaCl, 2x HALT protease inhibitor) and then purified on an Amylose resin column (NEB) and eluted using 10mM maltose diluted in lysis buffer. All proteins were concentrated on a 30 MWCO column (Amersham) into detergent-free buffer (1x PBS, pH 7.4, 5% glycerol) compatible for circular dichroism or IP LC-MS analyses. Proteins were also quantified via the Lowry Protein Assay (Creative Proteomics) and on a Coomassie-stained SDS-PAGE gel containing control concentrations of BSA (250 ng, 500 ng, and 1000 ng).

Circular dichroism was performed using a Jasco J-815 Spectrometer (OSU Biophysical Interaction & Characterization Facility; University of Pennsylvania Biological Chemistry Resources Center [BCRC]) as previously described [28]. In brief, 0.1 mg/mL of purified MBP-tagged proteins were analyzed in 0.1 mm width quartz cuvettes (Sigma), with scans taken every 1 nm from 190 to 250 nm at a controlled temperature of 4°C. CD values (mdeg) were corrected and converted to molar ellipticity in Excel (Microsoft) and values were plotted in Prism (GraphPad).

PG-conjugated beads and PA-conjugated beads were obtained from Echelon Biosciences (P-B0PG and P-B0PA, respectively). 0.9 nmol of purified MBP-tagged dSTART domain, its mutant versions (MBP-dLPSGFmut and MBP-dSDmut), or an MBP alone control were incubated with 0.25 nmols of beads equilibrated in Buffer System 1 (containing 0.3% IGEPAL). Mixtures were gently rotated at room temperature for 1 hour, then transferred to 4°C overnight for complete capture. Mixtures were gently washed three times using 500 µL of Buffer System 1, eluted in 1x Laemmli buffer, and separated from the beads after a 30 second spin at maximum speed. Bound proteins and 1:500 diluted inputs were separated on an SDS-PAGE gel. Proteins were detected via western blotting using a 1:5,000 dilution of HRP-conjugated antibody targeting the 6x-His tag at the C-terminus of all recombinant proteins (ThermoFisher).

### Liposome production, lipid immunoprecipitations, and mass spectrometry

Total lipids from 15 g of 12-day old *Arabidopsis* seedling tissues were extracted using the Bligh-Dyer extraction method [94]. Lipids were dried under argon and resuspended in 100% ethanol to 100 mg/mL. Liposomes were generated by rapid injection of these ethanol-suspended lipids into water. Liposomes were then diluted 1:10 into IP buffer (1x PBS pH 7.4, 150mM NaCl, 5% glycerol) and briefly sonicated. To identify lipids bound by the dSTART domain in plants, 100 mg of protein (mEGFP control and mEGFP-dSTART) was incubated with 6 mg of liposomes in IP buffer (1x PBS, pH 7.4, 150 mM NaCl, 5% glycerol) for 30 minutes at room temperature. Proteins were then re-purified using StrepTactin affinity chromatography, extensively washed to remove any liposome contamination, and eluted using 2.5 mM desthiobiotin. Lipids were then extracted using Bligh-Dyer and dried under argon. To identify lipids bound by the dSTART domain in bacteria, lipids were extracted from recombinant proteins immediately after purification from bacteria. Dried lipids were sent to Creative Proteomics for UPLC-MS analysis. A full list of identified lipids can be found in **Supp Table 3**.

Samples were extracted in 400 μL of isopropyl alcohol: MeOH (1:1, v/v), centrifuged for 15 min at 12,000 rpm at 4°C, then the supernatant was transferred for LC-MS analysis. Creative Proteomics uses an ACQUITY UPLC combined with AB SCIEX 5500 for ESI- and ESI+ modes. For ESI-, the LC system is comprised of ACQUITY UPLC BEH Amide (100×2.1mm×1.7 μm). Flow rate of the mobile phase is 0.1 mL/min, the column temperature is maintained at 45°C, and the sample manager temperature is set at 4°C. Mass spectrometry parameters in ESI- mode is listed as follows: Curtain Gas 35 arb, Collision GAS 9 arb, IonSpray voltage 4.5 KV, Temperature 450°C, IonSource Gas1 50 arb and IonSource Gas2 50 arb. For ESI+, the LC system is comprised of ACQUITY UPLC BEH C18 (100×2.1mm×1.7 μm). Flow rate of the mobile phase is 0.3 mL/min, the column temperature is maintained at 45°C, and the sample manager temperature is set at 4°C. Mass spectrometry parameters in ESI+ mode is listed as follows: Curtain Gas 35 arb, Collision GAS 7 arb, IonSpray voltage 5.2 KV, Temperature 450°C, IonSource Gas1 55 arb and IonSource Gas2 55 arb.

## ACKNOWLEDGEMENTS

We are grateful to Dr. Doris Wagner and members of the Wagner Lab for thoughtful comments that have improved the manuscript. We also thank Ryan Kubanoff of the Penn BCRC for technical assistance. Work in the Husbands Lab is funded by grants from the National Science Foundation (#2310356) and the National Institutes of Health (1R35GM158110) to AYH.

## SUPPLEMENTARY INFORMATION

**Supplemental Table 1 – Percent identity comparisons. a**, HD-ZIPIII paralogs. **b**, CNA orthologs. **c**, PDF2 orthologs. **d**, CNA orthologs (START to dSTART). **e**, Other StARkin domain classes.

**Supplemental Table 2 – ChIP-seq and RNA-seq related information. a**, Diffbind outputs of PHB, PHB-dΔ, and non-transgenic control. **b**, DEseq2 comparisons of 24h estradiol-induced *PHB\** lines versus mock. **c**, DEseq2 comparisons of 24h estradiol-induced *PHB-d*Δ*\** lines versus mock. **d**, ChIP-seq mapping statistics. **e**, RNA-seq mapping statistics.

**Supplemental Table 3 – Lipidomics datasets. a**, Negative ion species obtained for all GFP or GFP-dSTART samples. **b**, Positive ion species obtained for all GFP or GFP-dSTART samples. **c**, Subset of lipids enriched in LC-MS analyses of GFP-dSTART protein over GFP negative control proteins. Lipids were extracted immediately after purification from bacteria. **d**, Subset of lipids enriched in LC-MS analyses of GFP-dSTART protein over GFP negative control proteins. Lipids were extracted after plant liposome incubation and repurification.

**Supplemental Table 4 – Oligonucleotides used in study.**

