## Supplemental Figures for "A cryptic START domain regulates deeply conserved transcription factors"

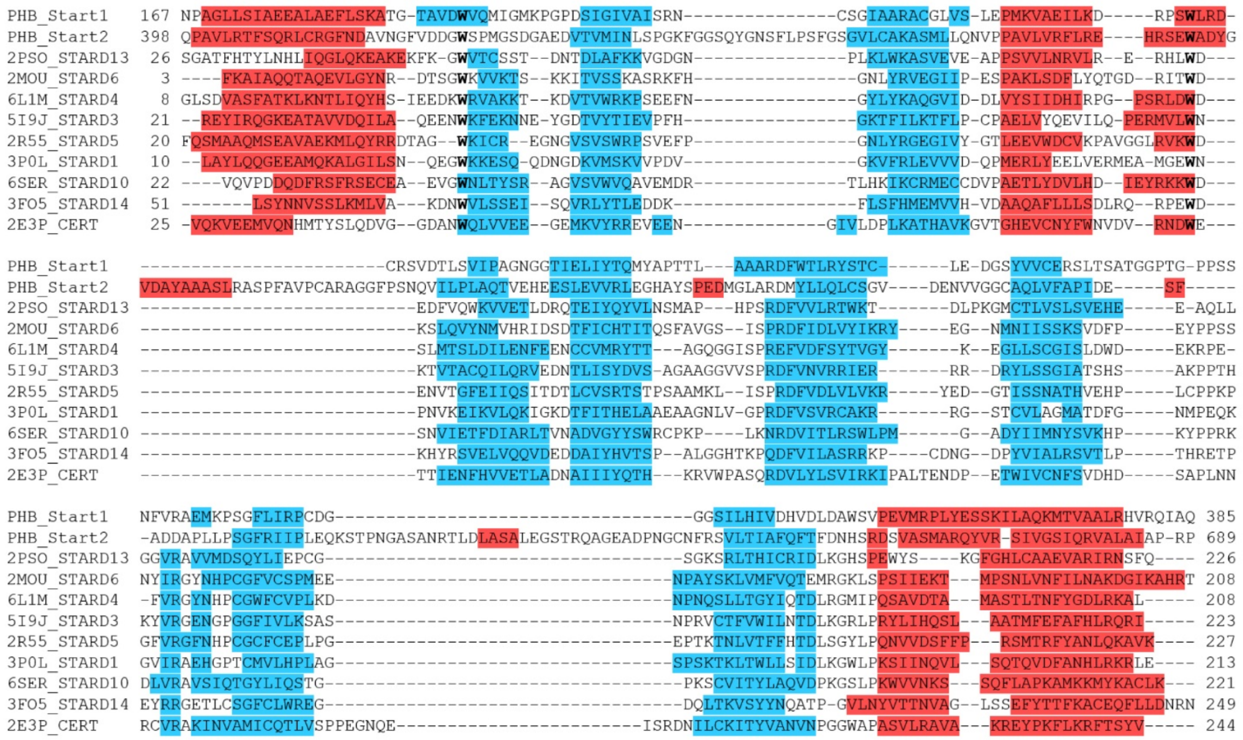

**Supplemental Figure 1. Multiple sequence alignment of uncharacterized region downstream of the START domain contains StARkin domain secondary structure features and patterns.** Secondary structure predictions for multiple StARkin domains made by Quick2D. Predicted α-helices are highlighted in red, and β-sheets are highlighted in blue. Bolded residues represent highly conserved tryptophan (W) residues that define StARkin domains. StARkin domains from the representative HD-ZIPIII TF PHB both share similar secondary structure patterns with mammalian START domains. Mammalian START domains are the top hits of the HHpred analysis.

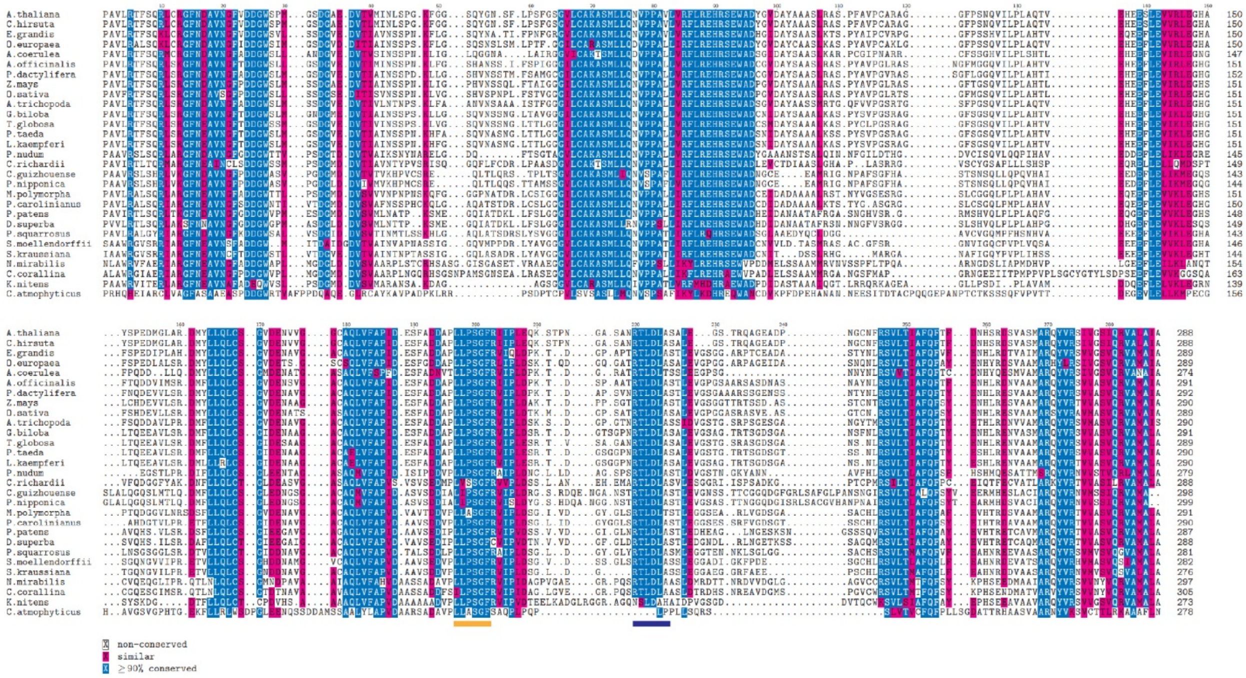


**Supplemental Figure 2. The ancestral-most HD-ZIPIII protein contains both StARkin domains.** Multiple sequence alignment of CNA orthologs reveals a deep conservation of the dSTART domain. The dSTART domain is present in the earliest-diverging organism known to contain an HD-ZIPIII protein, *Chlorokybus atmophyticus*. Regions of interest include the LPSGF (yellow bar) and RTLDLA (blue bar) motifs.

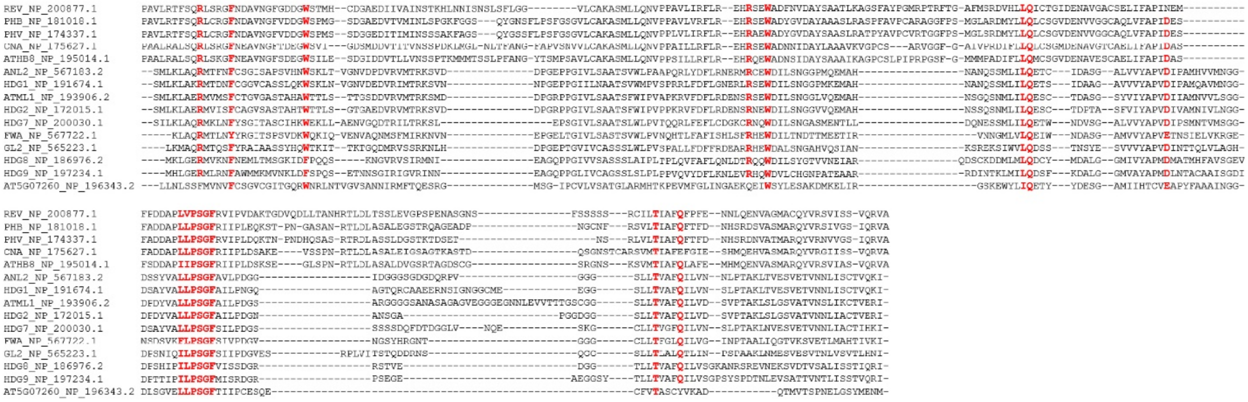

**Supplemental Figure 3. Multiple sequence alignment of dSTART domains from select *A. thaliana* HD-ZIPIII and HD-ZIPIV proteins.** Highly conserved residues (bold and red) are called from a multiple sequence alignment a consensus of more than 2800 HD-ZIPIII and HD-ZIPIV sequences (see **Supp Doc 1**). The dSTART domain is found in both HD-ZIPIII and HD-ZIPIV proteins, indicating a deep ancestry that predates the divergence of the two HD-ZIP subfamilies. The HD-ZIPIV dSTART domain also contains the highly conserved stretch of amino acids (LPSGF) that defines the dSTART domain.

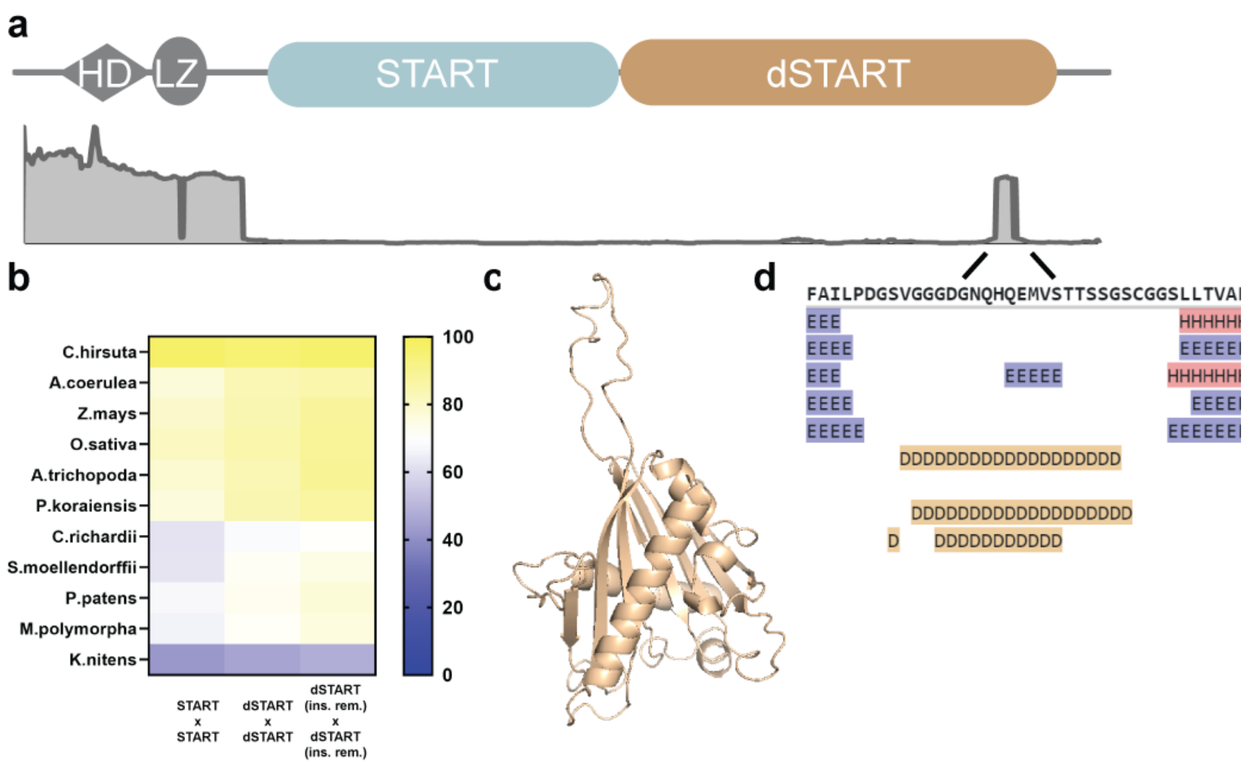


**Supplemental Figure 4. Structural and evolutionary analyses of a representative HD-ZIPIV family member PDF2. a**, PDF2 architecture (upper) with DEPICTER readout (lower; gray graph). Region near dSTART end also has predicted intrinsic disorder. **b**, Percent identity matrix comparing PDF2 START, dSTART, and dSTART (no insertions) sequences with orthologs in extant plant species showing an opposite pattern of conservation than HD-ZIPIII TFs. **c**, i-TASSER homology model of PDF2 dSTART domain with intrinsic disorder region (IDR) protruding as a large loop. **d**, Quick2D calls the IDR as disordered but does not predict any structures, in contrast to HD-ZIPIII TFs (**Fig. 1e**).

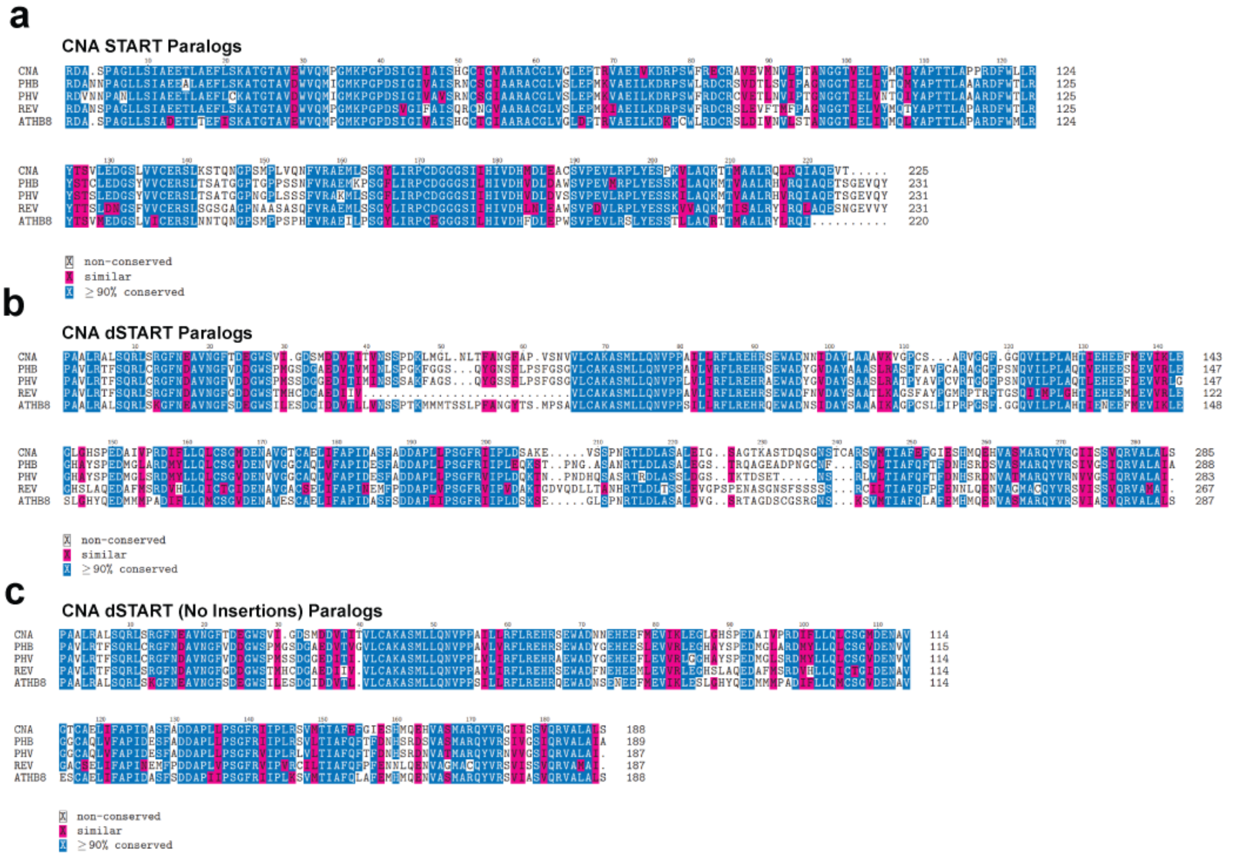


**Supplemental Figure 5. Multiple sequence alignments of START and dSTART domains in *A. thaliana* HD-ZIPIII paralogs. a,** Multiple sequence alignment showing conservation of HD-ZIPIII START domains. **b,** Multiple sequence alignment showing conservation of HD-ZIPIII dSTART domains. Note sequence conservation is lower for their insertions except at the RTLDLASXL stretch of the α3-β2 insertion. **c,** Multiple sequence alignment showing conservation of HD-ZIPIII dSTART domains after removal of the three dSTART-specific insertions.

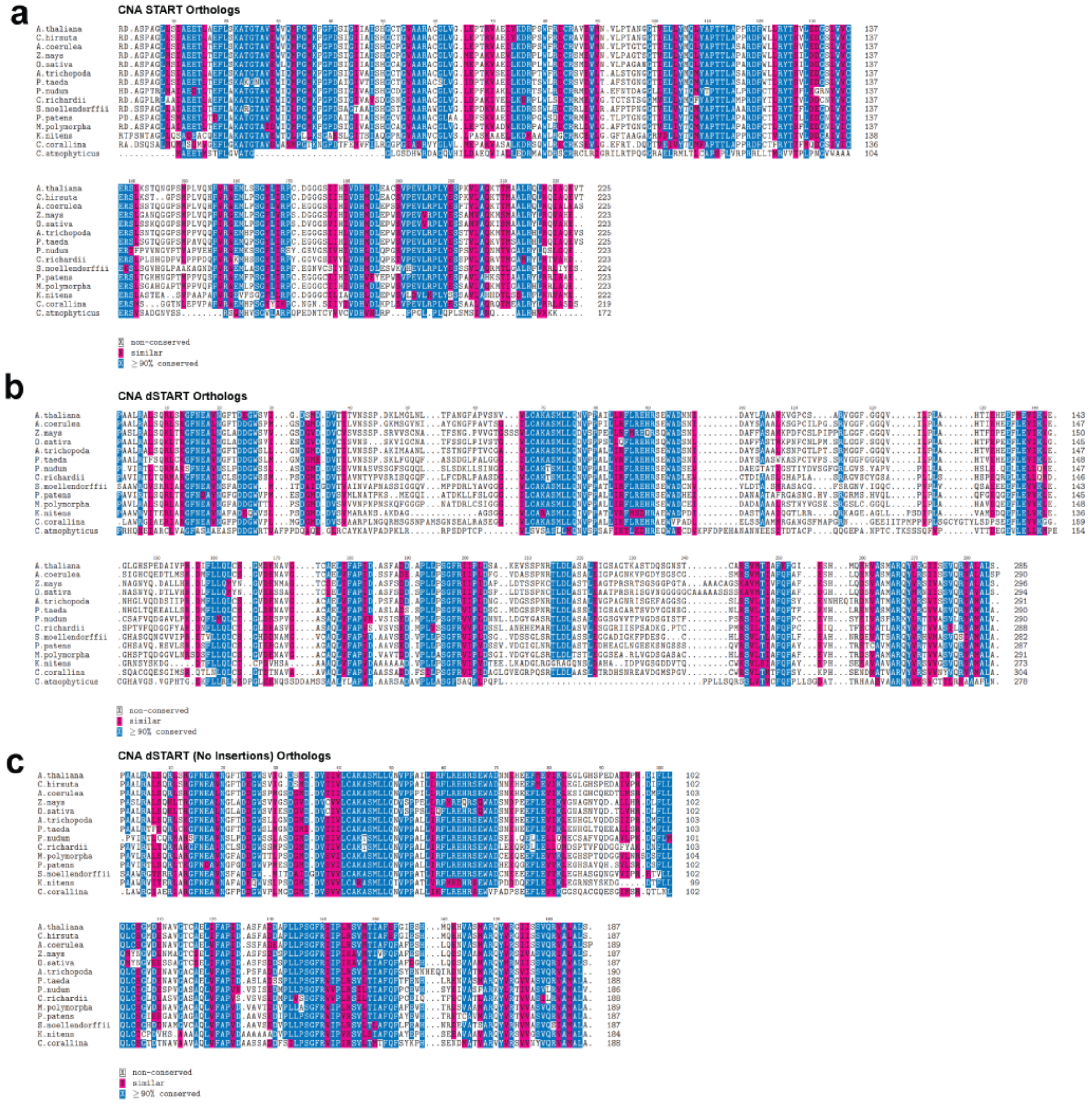


**Supplemental Figure 6. Multiple sequence alignments of START and dSTART domains from CNA orthologs across plant evolution. a,** Multiple sequence alignment showing conservation of CNA ortholog START domains. **b,** Multiple sequence alignment showing conservation of CNA ortholog dSTART domains. Note sequence conservation is lower for their insertions except at the RTLDLASXL stretch of the α3-β2 insertion. **c,** Multiple sequence alignment showing conservation of CNA ortholog dSTART domains after removal of the three dSTART-specific insertions.

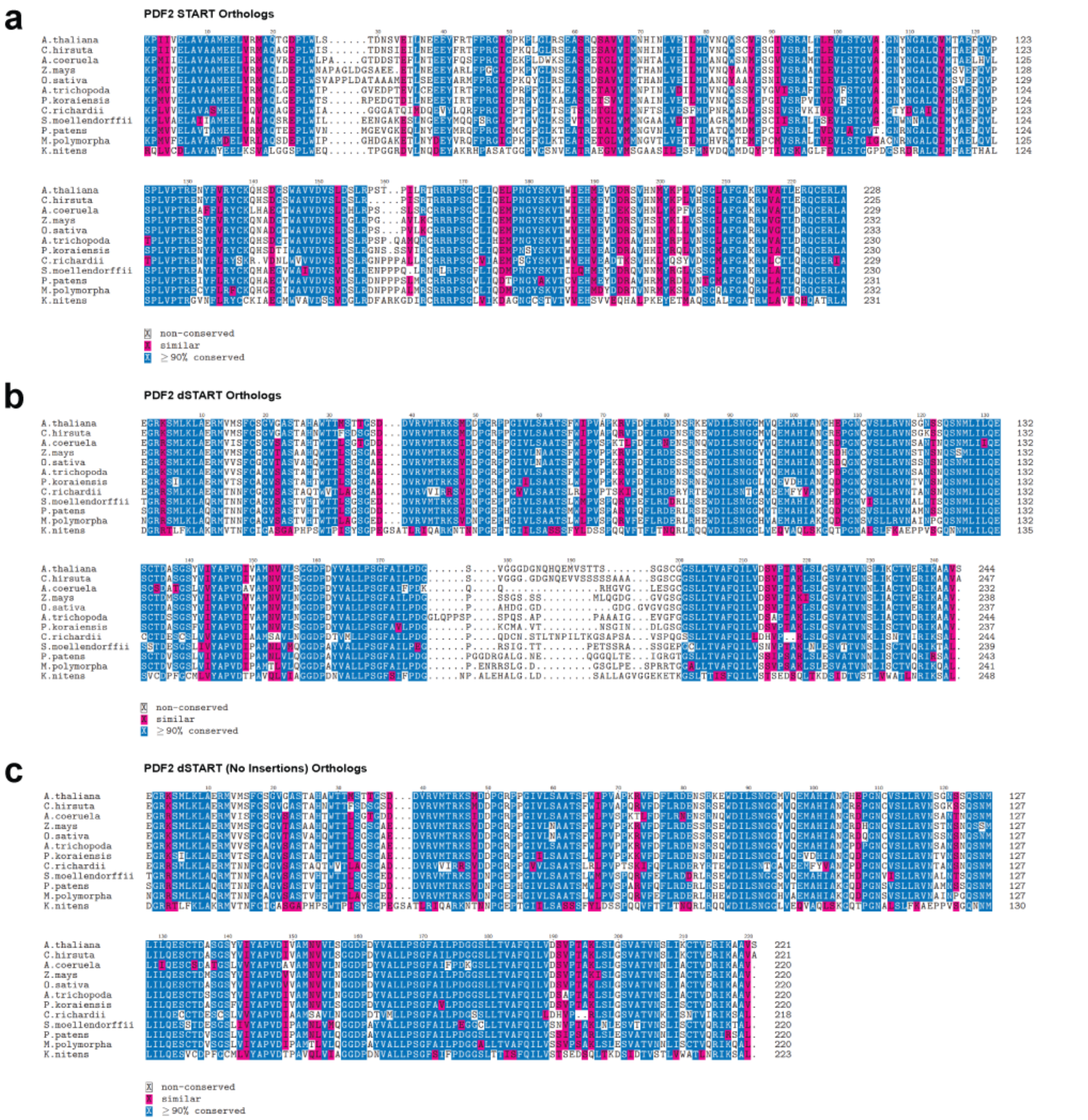


**Supplemental Figure 7. Multiple sequence alignments of START and dSTART domains from PDF2 orthologs across plant evolution. a,** Multiple sequence alignment showing conservation of PDF2 ortholog START domains. **b,** Multiple sequence alignment showing conservation of PDF2 ortholog dSTART domains. Note sequence conservation is lower for their insertions. **c,** Multiple sequence alignment showing conservation of PDF2 ortholog dSTART domains after removal of the three dSTART-specific insertions.

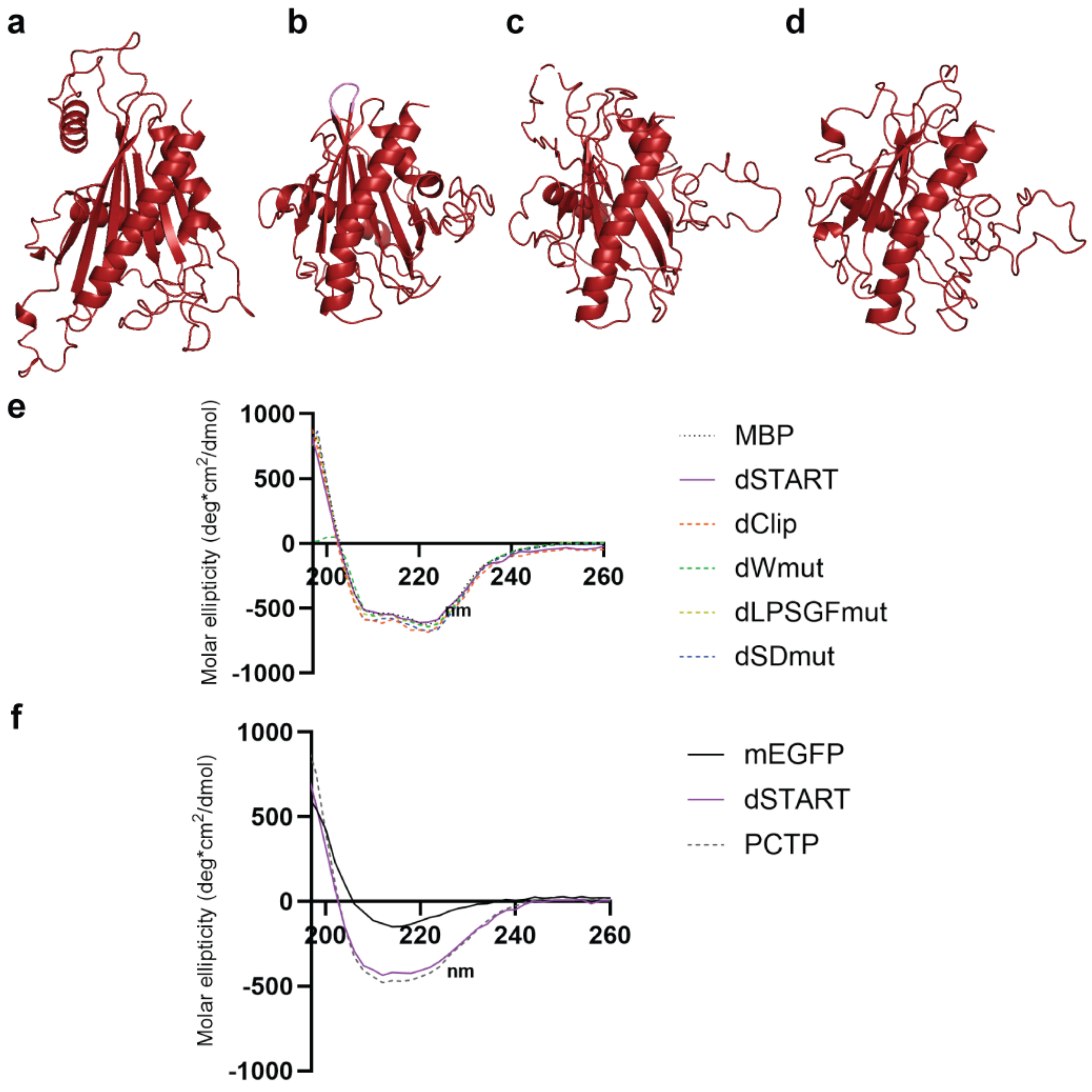


**Supplemental Figure 8. Homology models of CNA dSTART mutants.** **a**, W-to-A (dWmut; RMSD=0.667), **b**, clipped IDR (dClip; RMSD=1.011), **c**, SGF-to-TAY (dLPSGFmut; RMSD=1.864), and **d**, DAPLL and RDMYLLQ to AVVAA and GAVVGAG, respectively (dSDmut; RMSD=2.398). **e**, Circular dichroism of recombinant purified MBP-tagged START domains shows near identical secondary structures. Note the shoulder at ~215nm consistent with the numerous helices within these proteins. **f**, Circular dichroism of recombinant doubly-purified GFP-tagged START domains. GFP-dSTART and GFP-PC-TP have similar secondary structures that are distinct from GFP. Note the small shoulder at ~215nm in the START proteins consistent with the presence of their three $\alpha$-helices.

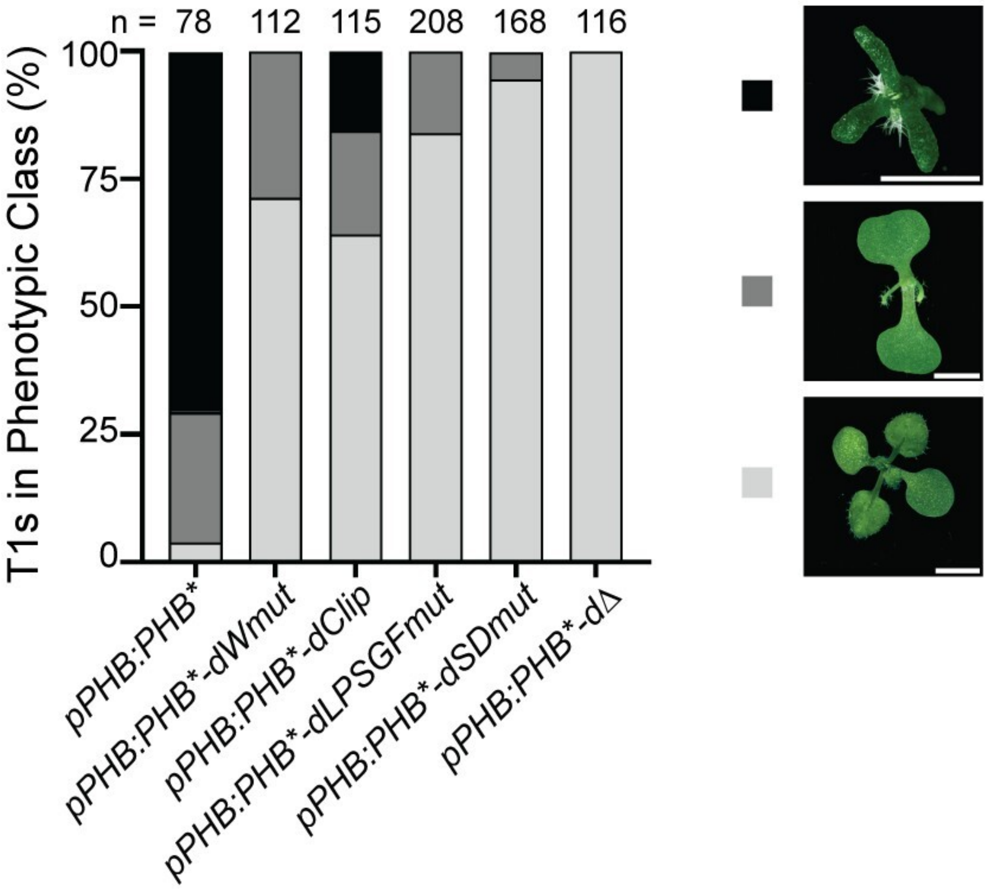


**Supplemental Figure 9. Phenotypes and scoring of primary transformants from modified *pPHB:PHB** reporter assay.** Phenotypic scoring of primary transformants (*n* above bar) carrying miR166-insensitive (*) constructs largely recapitulates *phb phv cna* complementation results. On the right are representative examples of severe (black), moderate (dark grey), and mild/wildtype (light grey) phenotypes. Categorization followed REF Husbands. Note bar graph is identical to graph in **Fig. 3b**.

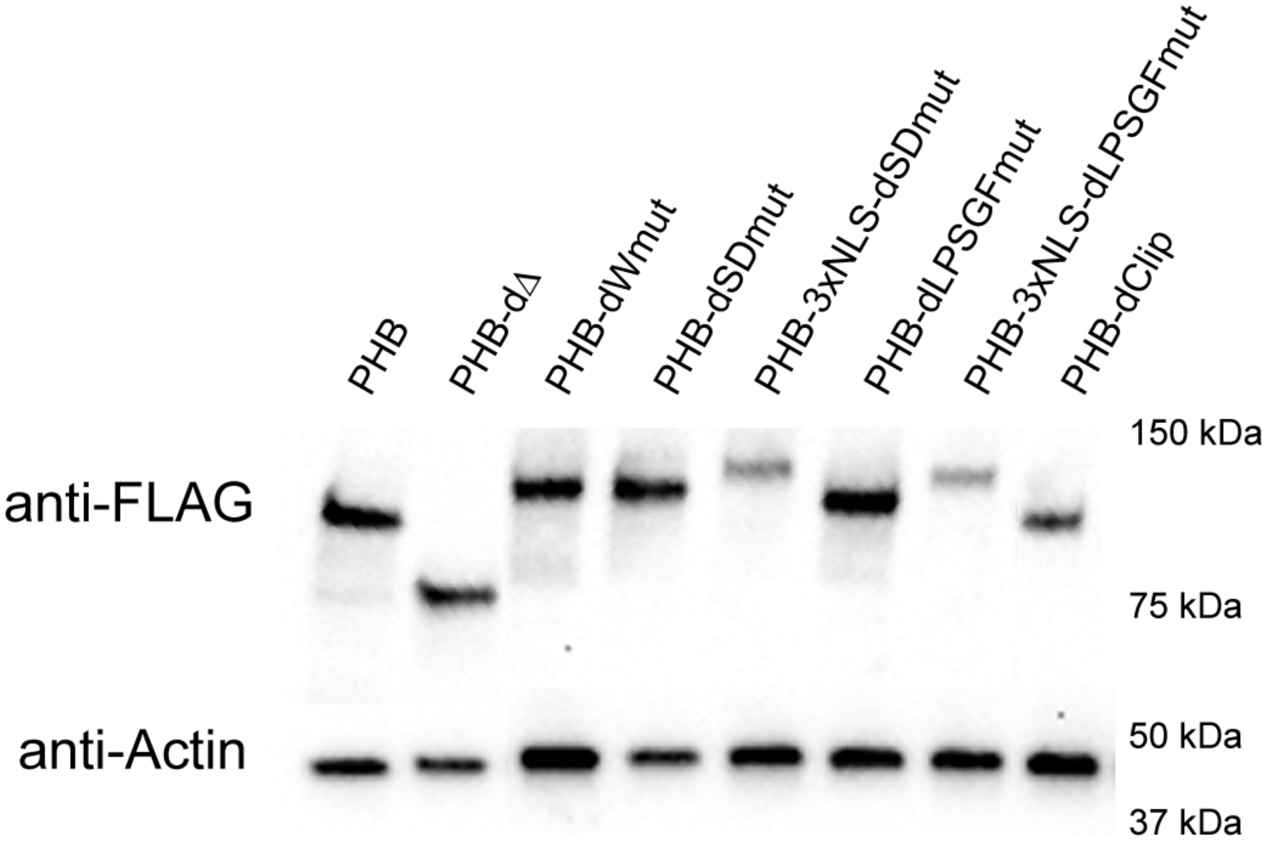


**Supplemental Figure 10. Protein stability is not affected by the dSTART domain.** Anti-FLAG Western blotting of estradiol-induced lines shows most PHB variants have no overt issues with protein accumulation. Two exceptions are 3xNLS-PHB-dLPSGFmut and 3x-NLS-PHB-dSDmut which accumulate to lower levels than their non-NLS-tagged counterparts.

**a**

| Protein Name | Position | Sequence | Score |
|---|---|---|---|
| >LocNES309912389_0 | 30-44 | MGSDGAEDVTVMINL | 0.033 |
| >LocNES309912389_0 | 61-75 | SFGSGVLCAKASMLL | 0.010 |
| >LocNES309912389_0 | 70-84 | KASMLLQNVPPAVLV | 0.003 |
| >LocNES309912389_0 | 72-86 | SMLLQNVPPAVLVRF | 0.014 |
| >LocNES309912389_0 | 100-114 | DAYAAASLRASPFAV | 0.004 |
| >LocNES309912389_0 | 150-164 | AYSPEDMGLARDMYL | 0.126 |
| >LocNES309912389_0 | 153-167 | PEDMGLARDMYLLQL | 0.536 |
| >LocNES309912389_0 | 190-204 | SFADDAPLLPSGFRI | 0.283 |
| >LocNES309912389_0 | 191-205 | FADDAPLLPSGFRII | 0.797 |
| >LocNES309912389_0 | 193-207 | DDAPLLPSGFRIIPL | 0.253 |
| >LocNES309912389_0 | 239-253 | DPNGCNFRSVLTIAF | 0.025 |
| >LocNES309912389_0 | 241-255 | NGCNFRSVLTIAFQF | 0.025 |
| >LocNES309912389_0 | 243-257 | CNFRSVLTIAFQFTF | 0.041 |
| >LocNES309912389_0 | 269-283 | ARQYVRSIVGSIQRV | 0.010 |
| >LocNES309912389_0 | 271-285 | QYVRSIVGSIQRVAL | 0.054 |

**b**

| Protein Name | Position | Sequence | Score |
|---|---|---|---|
| >LocNES488293312_0 | 30-44 | MGSDGAEDVTVMINL | 0.044 |
| >LocNES488293312_0 | 61-75 | SFGSGVLCAKASMLL | 0.010 |
| >LocNES488293312_0 | 70-84 | KASMLLQNVPPAVLV | 0.004 |
| >LocNES488293312_0 | 72-86 | SMLLQNVPPAVLVRF | 0.016 |
| >LocNES488293312_0 | 100-114 | DAYAAASLRASPFAV | 0.004 |
| >LocNES488293312_0 | 150-164 | AYSPEDMGLARDMYL | 0.126 |
| >LocNES488293312_0 | 153-167 | PEDMGLARDMYLLQL | 0.536 |
| >LocNES488293312_0 | 193-207 | DDAPLLPTAYRIIPL | 0.152 |
| >LocNES488293312_0 | 239-253 | DPNGCNFRSVLTIAF | 0.024 |
| >LocNES488293312_0 | 241-255 | NGCNFRSVLTIAFQF | 0.021 |
| >LocNES488293312_0 | 243-257 | CNFRSVLTIAFQFTF | 0.034 |
| >LocNES488293312_0 | 269-283 | ARQYVRSIVGSIQRV | 0.007 |
| >LocNES488293312_0 | 271-285 | QYVRSIVGSIQRVAL | 0.061 |

**c**

| Protein Name | Position | Sequence | Score |
|---|---|---|---|
| >LocNES901701744_0 | 30-44 | MGSDGAEDVTVMINL | 0.101 |
| >LocNES901701744_0 | 61-75 | SFGSGVLCAKASMLL | 0.010 |
| >LocNES901701744_0 | 70-84 | KASMLLQNVPPAVLV | 0.013 |
| >LocNES901701744_0 | 72-86 | SMLLQNVPPAVLVRF | 0.019 |
| >LocNES901701744_0 | 100-114 | DAYAAASLRASPFAV | 0.006 |
| >LocNES901701744_0 | 190-204 | SFADAVVAAPSGFRI | 0.088 |
| >LocNES901701744_0 | 191-205 | FADAVVAAPSGFRII | 0.424 |
| >LocNES901701744_0 | 193-207 | DAVVAAPSGFRIIPL | 0.115 |
| >LocNES901701744_0 | 239-253 | DPNGCNFRSVLTIAF | 0.025 |
| >LocNES901701744_0 | 241-255 | NGCNFRSVLTIAFQF | 0.022 |
| >LocNES901701744_0 | 243-257 | CNFRSVLTIAFQFTF | 0.041 |
| >LocNES901701744_0 | 269-283 | ARQYVRSIVGSIQRV | 0.007 |
| >LocNES901701744_0 | 271-285 | QYVRSIVGSIQRVAL | 0.061 |

**d**

| Protein Name | Position | Sequence | Score |
|---|---|---|---|
| >LocNES481266583_0 | 30-44 | MGSDGAEDVTVMINL | 0.101 |
| >LocNES481266583_0 | 61-75 | SFGSGVLCAKASMLL | 0.010 |
| >LocNES481266583_0 | 70-84 | KASMLLQNVPPAVLV | 0.017 |
| >LocNES481266583_0 | 72-86 | SMLLQNVPPAVLVRF | 0.025 |
| >LocNES481266583_0 | 100-114 | DAYAAASLRASPFAV | 0.015 |
| >LocNES481266583_0 | 180-194 | AQLVFAPIDESFAAV | 0.017 |
| >LocNES481266583_0 | 238-252 | DPNGCNFRSVLTIAF | 0.025 |
| >LocNES481266583_0 | 240-254 | NGCNFRSVLTIAFQF | 0.022 |
| >LocNES481266583_0 | 242-256 | CNFRSVLTIAFQFTF | 0.041 |
| >LocNES481266583_0 | 268-282 | ARQYVRSIVGSIQRV | 0.028 |
| >LocNES481266583_0 | 270-284 | QYVRSIVGSIQRVAL | 0.071 |

**Supplemental Figure 11. The dSTART domain contains three nuclear export signal (NES) sequences. a,** Amino acid stretches containing LSPGF (yellow) or SDmut (RDMYLLQ, DAPLL; blue) motifs emerge as top hits in the LocNES NES prediction software (LocNES REF). **b**, LPSGF, or **c**, SDmut mutations abolish their predictions as NES sequences. Note each set of mutations does not prevent the other motif(s) from being called as an NES suggesting independent contributions. **d**, dSTART proteins mutated for all three motifs show no NES and/or cytoplasmic retentions sequences.

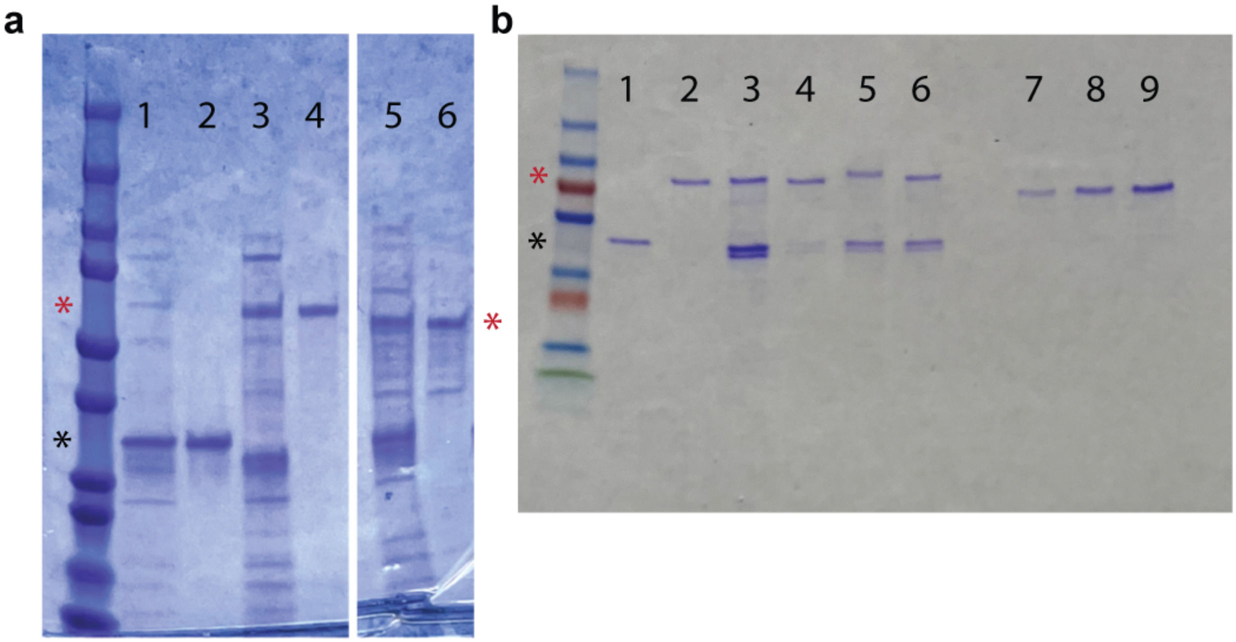


**Supplemental Figure 12. Recombinant StARkin domain proteins used in LC-MS and lipid-conjugated bead assays. a,** Double purification of GFP alone (lanes 1 and 2), GFP-tagged dSTART (lanes 3 and 4), and GFP-tagged PC-TP proteins (lanes 5 and 6) in *E coli*. Lanes 1, 3, and 5 are eluates from the first 6x-His-tag purification step using nickel affinity purification. Lanes 2, 4, and 6 are eluates from the second Twin-Strep-tag purification using Strep-Tactin XT. Note PC-TP samples were run on the same gel and intervening lanes were not included in the figure for clarity. **b**, Amylose resin-purified MBP alone (lane 1), MBP-dSTART (lane 2), MBP-dWmut (lane 3), MBP-dClip (lane 4), MBP-dLPSGFmut (lane 5), and MBP-dSDmut (lane 6). To assist with quantification, 250ng (lane 7), 500ng (lane 8), and 1ug (lane 9) of bovine serum albumin (BSA) was run out on the same gel. GFP or MBP alone negative controls are marked with a black asterisk. Approximate expected sizes of tagged StARkin variants are marked with a red asterisk.
